# SlMYB21 and SlMYB24 cooperatively regulate petal coloration and movement in *Solanum lycopersicum*

**DOI:** 10.64898/2026.10.19.760034

**Authors:** Moritz Friesch, Chen Zhang, Henrikje Smits, Julius Decker-Hibbel, Lena Harborth, Saskia Weinmar, Isabel Bühler, Antonio Santiago, Jone Echeverría, Frank Syrowatka, José Tomás Matus, Bettina Hause

## Abstract

In seed plants, reproductive success depends on the precise coordination of floral organ development. Here, we identify and characterize the *Solanum lycopersicum* transcription factor (TF) SlMYB24 as a member of the R2R3-MYB subgroup 19. While *Slmyb24* mutants exhibited no obvious phenotypic alterations, the previously characterized *Slmyb21* mutant displayed defects in floral development that were further exacerbated in the *Slmyb21 Slmyb24* double mutant, resulting in a severe phenotype characterized by female sterility and whitish petals that failed to open. Petals of *Slmyb21 Slmyb24* exhibited reduced xanthophyll content and elevated abscisic acid (ABA) levels, accompanied by reduced water content and decreased levels of primary sugars. Integration of transcriptome (RNA-sequencing) and cistrome (DNA affinity purification sequencing, DAP-seq) data identified genes involved in carotenoid and ABA biosynthesis and water transport as candidate regulatory targets of SlMYB21 and SlMYB24. These TFs activated the promoters of *PHYTOENE SYNTHASE2* (*PSY2*), *LYCOPENE β-CYCLASE (LCY-B*) and the petal-expressed aquaporin *TONOPLAST INTRINSIC PROTEIN1.1* (*TIP1.1*), whereas they repressed *9-*cis*-EPOXYCAROTENOID DIOXYGENASE1* (*NCED1*) promoter activity. In addition, SlMYB21 bound to the *SlMYB24* promoter and activated its expression. Promoter studies and electrophoretic mobility shift assays revealed that *SlMYB21* and *SlMYB24* bind the AC-rich DNA-binding motif ‘GTTAGGT’, which is to be found in the respective promoters and is consistent with the extensive overlap in their binding profiles. Together, these results reveal that the functions of SlMYB21 and SlMYB24 extend beyond fertility to coordinate petal pigmentation and opening.

## Introduction

Reproduction in seed plants depends on the formation of flowers containing highly specialized tissues that ultimately enable seed production and transmission of genetic information to the next generation. Successful reproduction requires a subset of morphological changes after flower meristem initiation. A typical flower consists of a concentric whorl of sepals, petals, stamens and carpels, with the latter two harboring the male and female structures (i.e., gametophytes), respectively. Petals are particularly the most prominent floral organs and display remarkable diversity in color, shape and size. During flower organogenesis, a tightly controlled genetic program is activated to ensure proper tissue specification and the subsequent development of sepals, petals, stamen and carpels (Endress, 2001). The tomato (*Solanum lycopersicum*) flower typically consists of five sepals, five partially fused petals, five fused stamen and two fused carpels, arranged in concentric whorls.

Petals not only provide a huge variety of colors but also play a crucial part in reproduction. In addition to providing visual cues for pollinators, petals can protect the reproductive tissues from abiotic stress (Karban et al., 2023). Petal development requires tightly coordinated cellular reprogramming, with cell proliferation predominating during early flower stages, followed by a phase of cell expansion during petal opening (Huang and Irish, 2015). Flower opening culminates in anthesis. When the flower becomes reproductively competent, pollen are released and pollination can occur, ultimately enabling fruit and seed development.

The reproductive fate of plants is regulated by signaling cascades triggered by phytohormones, including auxin, gibberellins (GA) and jasmonates (JA) (Browse, 2009; Pagnussat et al., 2009; Yamaguchi et al., 2014). The importance of JA is demonstrated by mutants impaired in JA biosynthesis or perception, which display fertility defects in both *Arabidopsis thaliana* and tomato (Li et al., 2001; Browse, 2009). In Arabidopsis, mutants of the JA-Ile receptor CORONATINE INSENSITIVE1 (COI1) or of JA-biosynthetic enzymes such as ALLENE OXIDE SYNTHASE (AOS) or OPDA-REDUCTASE3 (OPR3) are male sterile (Feys et al., 1994; Stintzi and Browse, 2000; Park et al., 2002). Transcriptomic analyses of JA-treated *Atopr3* stamens identified the R2R3-MYB transcription factors AtMYB21 and AtMYB24 as mediators of JA-regulated flower fertility (Mandaokar et al., 2006; Reeves et al., 2012). Accordingly*, Atmyb21* mutants exhibit a floral phenotype similar to that of JA-biosynthesis mutants, characterized by shortened filaments and non-viable pollen (Mandaokar et al., 2006). AtMYB21 interacts with its homologue AtMYB24, and the more severe phenotype of the *Atmyb21 Atmyb24* double mutant compared with *Atmyb21* indicates partially overlapping functions in the regulation of male fertility (Mandaokar et al., 2006; Song et al., 2013). In addition, overexpression of *AtMYB21* in *coi1* or *AtMYB24* in *opr3* partially rescued male fertility, further supporting central roles for both transcription factors (TFs) in stamen and pollen development (Song et al., 2011; Huang et al., 2017).

In tomato, the role of JA in fertility has been demonstrated in plants with defective JA signaling, such as *jasmonate-insensitive1* (*jai1*), and in mutants or transgenic lines impaired in JA biosynthesis, including *spr2*, *acx1a*, *SlAOC-*RNAi and *SlOPR3-*RNAi (Li et al., 2003; Li et al., 2005; Goetz et al., 2012; Bosch et al., 2014). Interestingly, these mutant and transgenic plants revealed a particularly important role for JA in female fertility, as indicated by the production of seedless fruits. Similar to AtMYB21, the tomato orthologue SlMYB21 is controlled by JA-signaling and regulates female fertility. *Slmyb21-2* plants exhibit normal stamen development but disrupted ovule development (Schubert et al., 2019). Female sterility results from premature cell death in the innermost cell layer of the nucellus, leading to interrupted ovule development and embryo abortion, and consequently parthenocarpic fruit formation (Schubert et al., 2019). Moreover, remarkable differences in the function of MYB21 TFs between Arabidopsis and tomato became obvious, since *Slmyb21* mutant flowers exhibit diminished JA levels signifying a positively regulatory role of SlMYB21 in regulating JA biosynthesis (Schubert et al., 2019).

Beyond reproductive organ development, subgroup 19 (S19) MYB transcription factors have been implicated in stress responses and specialized metabolism. AtMYB21 at least partially mediates JA-regulated wound response in *A. thaliana* (Mekkaoui et al., 2026). In grapevine (*Vitis vinifera*), VviMYB24 transcripts highly accumulate in the flower but also in the berry skin during ripening, directly regulating terpene and flavonol metabolism (Zhang et al., 2023). This broad control of specialized metabolism protects berries against ultraviolet and high-intensity visible light stress.

In many plants, including tomato, petal coloration is conferred by pigments such as carotenoids, which provide visual cues to attract pollinators (Lunau et al., 2018; Sinha et al., 2023). In addition to their roles as pigments, carotenoids are important secondary metabolites, which participate in photoprotection and serve as precursors for the biosynthesis of the phytohormones abscisic acid (ABA) and strigolactones (Demmig-Adams and Adams, 1996; Schwartz et al., 2003; Booker et al., 2005). Carotenoid biosynthesis in tomato has been studied predominantly in fruits, here lycopene accumulation confers the characteristic red color, and contributes to nutritional value (Clinton, 1998). Substrates of the carotenoid biosynthesis are isopentenyl diphosphate (IPP), and its isomer dimethylallyl diphosphate (DMAPP), which are converted to lycopene via geranylgeranyl diphosphate (GGPP) catalyzed among others by PHYTOENE SYNTHASE (PSY), PHYTOENE DESATURASE (PDS), and LYCOPENE-β-CYCLASE (LCY-B/CRTL-B) (for review see Liu et al., 2015). During fruit ripening, the MADS box transcription factor RIPENING INHIBITOR (RIN) regulates numerous ripening associated genes, including the carotenoid biosynthesis gene *PSY1* (Martel et al., 2011). In contrast to fruits, tomato petals mostly accumulate the yellowish xanthophylls, neoxanthin and violaxanthin, which together account for up to 90% of the total carotenoid content in petals and stamens (Isaacson et al., 2002).

The molecular regulation of petal movement has been predominantly investigated in horticultural species. In *Gentia scrapa*, diurnal petal bending depends on phosphorylation of plasma membrane aquaporin PLASMAMEMBRANE PROTEIN2.2 (GsPIP2.2) by the CALCIUM-DEPENDENT PROTEIN KINASE16 (Nemoto et al., 2022). Light-triggered phosphorylation activates GsPIP2.2, promoting water influx and swelling of cells on the adaxial side of the petal and thereby contributing to flower opening (Nemoto et al., 2022). Another mechanism involving asymmetric cell expansion has been described in *Rosa hybrida*. Here, the homeodomain transcription factor PETAL MOVEMENT-RELATED PROTEIN1 (RMP1), acting downstream of ethylene signaling via ETHYLENE INSENSITIVE3 (EIN3), activates endoreduplication in adaxial petal cells and also contributes to flower opening (Cheng et al., 2021).

Despite considerable progress in understanding the regulation of flower development in tomato, the transcriptional networks underlying specific processes, such as petal pigmentation and flower opening, and their relation to phytohormone signaling, remain poorly understood. Here, we investigate the overlapping and distinct functions of SlMYB21 and SlMYB24 in petal development and opening. While *Slmyb24* single mutants exhibited no obvious floral phenotype, simultaneous loss of SlMYB21 and SlMYB24 resulted in a substantially more severe phenotype than loss of SlMYB21 alone, characterized by frequent flower abortion and unopened, whitish petals with altered cellular morphology. To uncover the molecular basis of these phenotypes, we integrated transcriptomic profiling of *Slmyb21 Slmyb24* petals with DNA affinity purification sequencing (DAP-seq) binding profiles of SlMYB21 and SlMYB24. Our results identify both transcription factors as regulators of genes involved in carotenoid biosynthesis, ABA homeostasis and water transport, including the transcriptional activation of *TONOPLAST INTRINSIC PROTEIN1.1* (*TIP1.1*), encoding a tonoplast-located aquaporin, which we hypothesize regulates water movement in petals to ensure flower opening during the day. Moreover, we demonstrate that SlMYB21 directly binds to and activates the *SlMYB24* promoter, revealing a hierarchical relationship between the two S19-MYB transcription factors. Together, our findings expand the known functions of SlMYB21 and SlMYB24 beyond reproductive fertility and reveal their coordinated roles in petal pigmentation and opening.

## Material and Methods

### Plant material and growth conditions

Tomato seeds (*Solanum lycopersicum*) cv. MicroTom wild type, *Slmyb21-2* (Schubert et al., 2019), and mutants generated during this work were germinated on expanded clay for 10 days and seedlings then transferred to soil. Growth of tomato plants was done in a growth chamber with 16 h light (200 mmol photons*m^-2^*s^-1^; Metal Halide LampMT250DL, IWASAKI Electric Co., www.eye.co.jp) at 26°C and 8 h dark at 20°C, both at 70% humidity, plants for seed propagation were grown in the greenhouse. *Nicotiana benthamiana* plants were grown under a 16h light/ 8h dark photoperiod at a constant temperature of 24°C. The *Slmyb21-2* and *Slmyb24-3* (see below) mutants were crossed to obtain *Slmyb21-2 Slmyb24-3* heterozygous plants in F1. After self-pollination, F2 plants were screened for *Slmyb21* and *Slmyb24* homozygosity. For multiplication, plants homozygous for *Slmyb24* and heterozygous for *Slmyb21* were used. Homozygous double mutants were then selected for *Slmyb21* homozygosity by PCR and subsequent Sanger Sequencing of the resulting DNA fragment according to Schubert et al. (2019). Developmental stages of flowers were determined according to Dobritzsch et al. (2015).

### Phylogenetic analysis

The phylogenetic tree was conducted using S19-SlMYB (SlMYB21 and SlMYB24) protein sequences and those from previously characterized S19-MYB in 9 plant species, including *Arabidopsis thaliana* (AtMYB21, AtMYB24, AtMYB57) (Cheng et al., 2021), *Nicotiana langsdorffii X N. sanderae* (LxS-MYB305) (Liu et al., 2009), *Antirrhinum majus* (AmMYB305) (Liu et al., 2009), *Petunia x hybrida* (PhEOB1 and PhEOB2) (Spitzer-Rimon et al., 2010; Spitzer-Rimon et al., 2012), *Petunia axillaris* (PaEOB1 and PaEOB2) (Chopy et al., 2023), *Malus domestica* (MdMYB305) (Kurilla et al., 2019), *Fragaria x ananassa* (FaEOB2) (Medina-Puche et al., 2015), and *Vitis vinifera* (VviMYB24) (Zhang et al., 2023). Multiple sequence alignment of S19-MYB protein sequences was performed using MAFFT (v7.526) with the ‘--auto’ option. Maximum-likelihood phylogenetic analysis was conducted based on the aligned protein sequences using IQ-TREE (v2.4.0), and node support was evaluated using 1000 bootstrap replicates (−bb 1000). The phylogenetic tree was visualized by FigTree (v1.4.4).

### Subcellular localization in *N. benthamiana*

The CDS of *SlMYB24* was amplified using gene-specific primers with matching overhangs for level 2 GoldenGate assembly and cloned in frame with the CDS of *GFP* (Werner et al., 2012). *N. benthamiana* protoplasts were transformed via PEG mediated transformation according to Schubert et al. (2019). After overnight incubation, subcellular localization was investigated using a LSM900 (Zeiss GmbH, www.zeiss.de). Fluorescence of GFP and chlorophyll was recorded upon excitation with 488 nm and emission at 505-550 nm and at 644-800 nm, respectively.

### Generation of *Slmyb24* mutants

To create a CRISPR/CAS9 construct for editing *SlMYB24*, two guide RNAs were designed according to Crispr-P 2.0 (Lei et al., 2014), crispr.hzau.edu.cn/CRISPR2/). The guide RNAs were then cloned in-frame with two U6 RNA promoters into a binary vector containing the CAS9 cassette. Transformation was done using cotyledons of 10-day-old seedlings according to Schubert et al. (2019). After successful regeneration of plants, they were screened for deletions at the *SlMYB24* locus with gene specific primers (Supplementary Table S1). Homozygous plants were obtained in the F2 generation.

### Time series of flower opening

Twenty pictures per hour were taken from 45 days old plants over several days using a Canon 700D camera (www.canon.de). A digital intervalometer (Magic Lantern, www.magiclantern.fm) was used for this purpose. The individual photos were compiled into a video using the FFmpeg git-b92577405b program (www.ffmpeg.org). One second in the video corresponds to one hour, i.e. 20 frames per second. The videos were edited using Shotcut Portable 25 and DaVinci Resolve 19.1.4 (www.shotcut.org).

### DNA affinity purification sequencing (DAP-seq)

The coding sequences of *SlMYB21* and *SlMYB24* were amplified and cloned into the pIX-HALO expression vector, generating N-terminal HALO-tagged fusion proteins. Genomic DNA was extracted from young leaves of tomato (*Solanum lycopersicum* cv. MicroTom). DAP-seq library preparation, sequencing, and downstream bioinformatic analyses were performed largely following previously established procedures (Zhang et al., 2023) and the standardized protocols and computational pipelines described in (Zhang et al., 2026). Sequencing reads were mapped to the tomato Heinz reference genome assembly SL4.0. For *de novo* DNA-binding motif discovery, 200-bp genomic sequences centered on the binding events identified by GEM were retrieved from the 600 most enriched peaks for each TF. Motif enrichment analysis was subsequently performed using STREME, implemented in MEME Suite v5.5.3 (www.meme-suite.org/meme/), with default parameters unless otherwise specified.

### Transcript profiling

Petals were harvested from WT and *Slmyb21 Slmyb24* flowers at stage 3 and stage 6, flash frozen and homogenized with mortar and pistil under liquid nitrogen. Total RNA was isolated using the Innu Plant prep Kit (InnuStar, https://innuscreen.ist-ag.com). Quality and integrity of RNA were assessed using 1% Agarose gel. RNA from three biological replicates per genotype and developmental stage were submitted to Novogene (https://eu.novogene.com) for mRNA paired-end (150 bp) sequencing on a NovaSeq6000 System.

The Illumina raw sequence reads were trimmed using fastq version 2.0. The minimum PHRED score accepted for trimming was 20 and reads shorter than 40 bp were discarded. Reads were aligned against the reference Heinz SL4.0 genome assembly using the HISAT2 v2.1.0 aligner with default parameters. Aligned reads were counted with Feature Counts, using the previous *in-house* merged ITAG4.0+4.1 annotation. A differential expression analysis was performed for each developmental stage separately, comparing mutant with the WT using R package limma. The counts of aligned reads were normalized with the TMM method implemented in the edgeR package. Genes that showed a minimal log2-fold-change of ±1 and had an adjusted p-value ≤ 0.05 as calculated by the Benjamini-Hochberg method were considered as differentially expressed genes.

### Gene expression analysis

Total RNA was isolated from homogenized frozen tissue with the Innu Plants RNA prep Kit (InnuStar) according to the manufacturer’s instructions and treated with the DNA free DNA Removal Kit (Thermo Fisher, www.thermofisher.com). First strand cDNA synthesis from 1 µg DNA-free RNA was carried out using the Revert Aid H-First strand Polymerase Kit (Thermo Fisher) using an equal molar mix of random and poly(T) oligonucleotides. Real-time quantification of gene expression was performed at least with 3 biological replicates with CDS specific primers (Supplementary Table S1) using the qPCR Mastermix EvaGreen® No Rox (Bio&Sell, www.bio-sell.de). Relative transcript levels were calculated using delta-CT method and normalized to *TIP41* (Solyc10g049850) (Expósito-Rodríguez et al., 2008). Primers are given in Supplementary Table S1.

### Transcriptional activation analysis in yeast cells and yeast-two hybrid assay

To test the transcriptional activity of SlMYB24, its CDS was fused to GAL4BD into pDEST32. Yeast transformation was performed as described (Schubert et al., 2019). Transcriptional activity was deduced from yeast growth on synthetic defined (SD) medium Trp/-Leu/-Ala or -Trp/-Leu/-His (+ 2 mM 3-amino-1,2,4-triazol).

For yeast-two-hybrid assays, transformation of competent pJ69 yeast cells was done as described previously (Gietz and Schiestl, 2007; Schubert et al., 2019). To prevent self-activation, the activation domain of SlMYB21 was deleted. Yeast cells were grown for 6 d on SD medium -Trp/-Leu. Single colonies were selected and cultured overnight at 30°C to an OD600 of 1, harvested, resuspended and dotted either on SD medium - Trp/-Leu/-Ala or on SD medium -Trp/-Leu/-His (+ 2 mM 3-amino-1,2,4-triazol) and incubated at 30°C for 3 d. Cells grown on SD medium -Trp/-Leu/-His were indicative for an interaction of the proteins under study.

### Bimolecular fluorescence complementation (BiFC) and split-luciferase (Luc) assays

BiFC assays were conducted using 2in1-vectors (Grefen and Blatt, 2012). Transformation of *N. benthamiana* protoplasts and imaging was done according to Schubert et al. (2019).

For split-Luc-assays, the full-length CDS of *SlMYB21* and *SlMYB24* were either cloned in frame into the pCambia1300-cLuc or the pCambia1300-nLuc plasmids using the OneStep In Fusion Kit (Takara, ww.takarabio.com) to generate the binary expression constructs. The resulting expression constructs were transformed into *Agrobacterium tumefaciens*, which were then used in a 1:1 mixture to infiltrate *N. benthamiana* leaves. Three days after infiltration, leaves were harvested and flooded for three minutes with substrate mixture (1.07 mM MgCO₃, 20 mM Tricine, 0.1 mM EDTA, 2.67 mM MgSO_4_ꞏH_2_O, 135 mM Coenzym A, 200 mM ATP, 10 mM DTT, 1 mM Luciferin). Three minutes after quenching in the dark, light signals were detected using the Fusion system (Vilber, www.vilber.com). Pictures are shown in false coloration. Experiments were done 3 times with similar results.

### Histochemical Analyses

Immediately after harvesting, petals were trimmed to 4 mm from the base of the petal in the fixative and embedded in polyethylene glycol (PEG) according to Dobritzsch et al. (2015). The embedded petals were sectioned into 7–10 µm thick sections, which were transferred to a poly-*L*-lysine-coated slide and stained with Calcofluor White and Basic Fuchsin according to Ursache et al. (2018). The sections were analyzed using an AxioImager microscope (Zeiss) with excitation/emission of 380 nm/425–465 nm and 532 nm/600–650 nm for Calcofluor White and Basic Fuchsin, respectively, and photographed using an Axiocam MRc5 (Zeiss). The surface area of the epidermal cells within the cross-section were measured using Fiji (ImageJ v1.54).

Scanning electron micrographs of fresh, unfixed petals were made with an ESEM XL-30 FEG (FEI/Philips, www.thermofisher.com) operating in the wet mode. The gas pressure in the ESEM chamber (1.5 mBar) was regulated by introducing water vapor. A gaseous secondary electron detector was used for imaging.

### Measurement of carotenoid, jasmonate and abscisic acid contents

Petals for carotenoid measurements were harvested and ground under liquid nitrogen. Carotenoids were extracted with 96 % ethanol after adding 500 μl of ethanol to approximately 70 mg of frozen powder, mixing and centrifugation at 3 °C and 13,000 rpm for five minutes (Moros et al., 2002). This was repeated three times with 500 μl of ethanol and once with 200 μl of ethanol, after which all pellets were completely decolorized. The HPLC analysis of combined ethanol extracts was carried out using the Waters Alliance® e2695 Separation Module (Waters Corporation, www.waters.com) using the EC HPLC column m(analytical), NUCLEODUR^®^ (MACHEREY-NAGEL GmbH & Co., www.mn-net.com). Absorbance was measured from 240 nm to 600 nm and evaluated at 445 nm using the 2998 PDA Detector (Waters Corporation).

JA-Ile and ABA were extracted from 20-50 mg of fresh material according to Balcke et al. (2012). Isotope-labelled compounds ([^2^H_6_]JA, [^2^H_2_] JA-Ile and ^2^H_6_-ABA, 50 ng each) were added as internal standards. Hormone contents were determined after separation in the ACQUITY UPLC System (Waters Cooperation) and detection by ESI-tandem mass spectrometry (MS/MS) using 3200 Q TRAPR LC/MS/MS mass spectrometer (Waters Cooperation).

### Measurement of water and sugar content

To determine the water content, three petals were harvested from five individual plants of each genotype and combined into five pools. The fresh weight (FW) was determined immediately after collection. The samples were then dried at 50 °C for one week, and the dry weight (DW) was determined. For sugar content determination, petals at developmental stages 3 and 6 were used. Measurements of glucose, fructose and saccharose were performed according to Schaarschmidt et al. (2004).

### *Trans*-activation assay

To obtain effector constructs, cDNAs of *SlMYB21* and *SlMYB24* were amplified using the GXL Polymerase and specific primers (Supplementary Table S1). The resulting fragments were cloned into the pDGB3_alpha vector using the ClonExpress II One Step Cloning Kit (Takara) according to the manufacturer’s protocol. Positive constructs were introduced into *A. tumefaciens* strain GV1301. The used promoter regions upstream of ATG (*upr*) were cloned in the pCambia1302-Luc vector upstream of the ATG of *luciferase*. *Trans*-activation assays were performed as described previously (Panda et al., 2022; Man et al., 2026). In brief, *A. thumefaciens* strains containing effector and reporter constructs were used for co-infiltration of *N. benthamiana* with a ratio of 4.5:1 (effector:promoter). For co-infiltration of two transcription factors the ratio was 4.5:4.5:1. After three days, leaf material was harvested, flash frozen and grinded under liquid nitrogen. Measurements were conducted using the microplate reader Spark (Tecan, www.tecan.com). *Trans*-activation assays were performed with at least six biological replicates, and luciferase activity was normalized to renillase activity.

### Electrophoretic mobility shift assay (EMSA)

The SlMYB21 and SlMYB24 CDS was amplified by PCR and introduced to the pHLT plasmid containing the sequence for an N-terminal 6xHis-Lypoyl domain (Golovenko et al., 2018). Using the *E. coli* Rosetta strain, expression of *6xHis-Lipoyl-SlMYB21* or *6xHis-Lipoyl-SlMYB24* was induced by addition of 1 mM IPTG at 19°C overnight. Fusion protein was purified using a HisTrap HP column (Cytiva, www.cytivalifesciences.com). Purified His-Lyp-SlMYB21 and His-Lyp-SlMYB24 were incubated with Cy5-probe, either alone or with WT-competitor or mut-competitor at room temperature for 20 min. Sequences of probes are given in Supplementary Table S1. EMSA samples were run on a 4% polyacryl amide gel at 100 V for 1 h. Cy5 fluorescence at 665 nm was recorded upon excitation with 635 nm on a Typhoon FLA 9500 scanner (General Electric, www.ge.com).

### Statistical analysis

Statistical analyses were performed using GraphPad Prism (www.graphpad.com) and applied to the different datasets as indicated in the figures. PHOTOSHOP 12.0.4 (Adobe Systems, http://www.adobe.com) was used to process micrographs and photographs.

## Results

### SlMYB24 is a flower specific transcription factor of R2R3-MYB subgroup 19

In *A. thaliana*, it is known that subgroup 19 (S19) members of R2R3-MYB, including AtMYB21, AtMYB24, and AtMYB57, act downstream of JA, interact and regulate filament elongation and pollen maturation (Reeves et al., 2012; Cheng et al., 2021). This encouraged us to investigate whether S19-MYBs in *S. lycopersicum* are similarly involved in flower development. In *S. lycopersicum*, R2R3-MYB transcription factor (TF) family has been previously reported to contain only one S19 member, *SlMYB21* (Solyc02g067760) (Gates et al., 2016). SlMYB21 was subsequently functionally characterized as a regulator of flower and ovule development as well as a positive regulator of jasmonate biosynthesis providing a positive feed-back in JA/JA-Ile accumulation during early stages of flower development (Niwa et al., 2018; Schubert et al., 2019). Here, we identified *SlMYB24* (Solyc02g086690) as a previously unrecognized S19 member and the closest homologue of *SlMYB21*. Due to a wrong structural annotation of *SlMYB24*, this gene was not classified as S19 in the characterization of R2R3-MYB TF family. Its exon/intron structure was incorrectly annotated, with the second intron prematurely interrupted and the incorrect coordinates of the third exon (Supplementary Figure S1A). The typical motif of S19 in the C-terminus was missing in the original coding sequences of *SlMYB24*. *SlMYB24* was therefore reannotated to align with the available transcriptomic data.

Phylogenetic analysis confirmed that *SlMYB24* is an ortholog of *AtMYB24* and belongs to R2R3-MYB subgroup 19 (Supplementary Figure S1B). To assert in which organs *SlMYB24* may have a regulatory role, and to compare its gene expression boundaries with *SlMYB21*, two public platforms, Tomato eFP Browser (https://bar.utoronto.ca/efp2/Tomato/Tomato_eFPBrowser2.html) and TomExpress (Zouine et al., 2017), were explored. We found that *SlMYB24* transcripts are enriched in flower buds and reached its peak in open flowers while being absent in seeds, fruits, and vegetative tissues (Figure 1A). By dissecting flower organs, expression analysis showed that *SlMYB24* transcript levels increased gradually in petals, stamens, and carpels from developmental stage 1 to stage 6, with peak expression observed at stage 6 in all three organs (Figure 1B). As SlMYB24 is classified as a R2R3 transcription factor, we confirmed its transcription activity by yeast one-hybrid assay (Figure 1C). Subcellular localization studies in *N. benthamiana* protoplasts confirmed the nuclear localization of SlMYB24 (Figure 1D).

**Figure 1.**
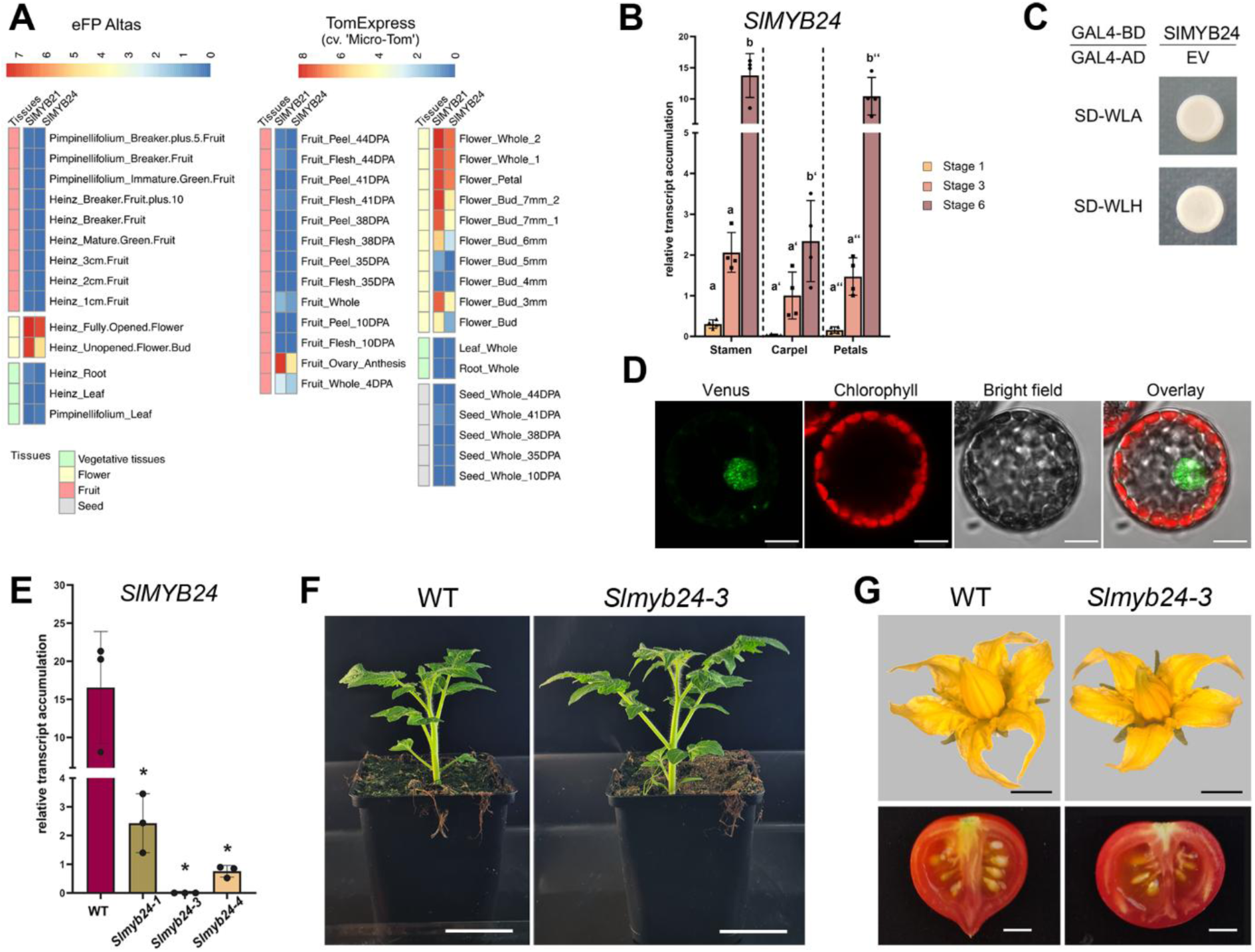
SlMYB24 is a flower specific transcription factor of the R2R3-MYB subgroup 19. A) *SlMYB24* transcripts are predominantly detected in flower buds and open flowers, as revealed by expression data retrieved from the tomato eFP browser (http://bar.utoronto.ca/efp_tomato/cgi-bin/efpWeb.cgi) and the TomExpress database (Zouine et al., 2017). B) Expression profiles of *SlMYB24* in different floral organs during flower development. Stamen, carpels and petals were collected at developmental stages 1, 3 and 6. Transcript abundance was determined by RT-qPCR and normalized to the reference gene *TIP41*. Data represent means ± SD (n=4 biological replicates, each comprising three pooled organs). Different letters indicate statistically significant differences within each flower organ, as determined by two-way ANOVA followed by Tukey’s post hoc test (P≤0.05). C) SlMYB24 functions as a transcriptional activator in yeast. Yeast cells transformed with constructs encoding SlMYB24 fused to GAL4-DNA-binding domain (BD) and the GAL4 activation domain (AD) only (empty vector) were grown on SD medium without Trp, Leu and Ala (-WLA) or without Trp, Leu and His (-WLH). Growth on these media confirmed successful transformation and transcriptional activity of SlMYB24. D) Subcellular localization of SlMYB24 in *Nicotiana benthamiana* mesophyll protoplasts transiently expressing the *35Spro:gSlMYB24* construct. Green fluorescence indicates nuclear localization of the SlMYB24–VENUS fusion protein, while red fluorescence corresponds to chloroplast autofluorescence. Scale bars = 10 µm. E) *Slmyb24-3* has no detectable *SlMYB24* expression. Expression of *SlMYB24* in the wild type and different CRISPR/Cas9 knockout alleles was measured in carpels of flowers at stage 6 by RT-qPCR and set in relation to *TIP41*. Mean values ±SD are shown (n = 3). Data were compared between wild type and *Slmyb24* alleles by one-way ANOVA with Dunnett’s multiple-comparison test (\**P* ≤ 0.05). F) Representative photographs of five-week-old WT and *Slmyb24-3* plants showing no obvious differences in vegetative development. Scale bars = 5 cm G) Representative photographs of WT and *Slmyb24-3* flowers and fruits, showing normal floral morphology and seed-containing fruit development in the mutant. Scale bars = 5 mm. Supporting data: Supplementary Figures S1-S3

Our previous work showed female sterility in *Slmyb21* mutants (Schubert et al., 2019). To investigate the contribution of SlMYB24 to flower development and fertility as well as its potential redundancy with SlMYB21, we generated CRISPR/CAS9-mediated knock-out mutants of *SlMYB24*. Screening of F2 generation revealed three mutant alleles. Out of these three alleles, line *Slmyb24-3* showed no detectable *SlMYB24* transcripts (Figure 1E). The *Slmyb24-3* allele carries a 7 bp deletion at position 56-62 of exon 1 resulting in a premature stop codon (Supplementary Figure S2). Plants of *Slmyb24-3* (hereafter called *Slmyb24*) showed no altered vegetative and flower phenotype compared to the wild type (Figure 1F, G). Even the transcript levels of *SlMYB21* were not altered in carpels and petals of these mutants in comparison to wild type (Supplementary Figure S3). Accordingly, mutants produced normal fruits which contained fertile seeds as the wild type (Figure 1G). This indicated that SlMYB24 might be indispensable for proper flower and seed development.

AtMYB21 and AtMYB24 are known to interact physically via their C-terminal activation domain in *A. thaliana* (Song et al., 2011), and At*myb21 Atmyb24* double mutant exhibits a strong floral phenotype, indicated by disrupted male fertility as well as smaller petals (Cheng et al., 2009; Reeves et al., 2012). To test, whether SlMYB21 and SlMYB24 do interact, yeast two-hybrid (Y2H), bimolecular fluorescence complementation (BiFC), and split-luciferase (Split-Luc) assays were performed and confirmed their interaction *in vivo* (Figure 2A-C). We then crossed *Slmyb21-2* mutant (hereafter called *Slmyb21*) with *Slmyb24* mutant to obtain the double mutant *Slmyb21-2 Slmyb24-3* (hereafter called *Slmyb21 Slmyb24*) to assess the genetic interaction between SlMYB21 and SlMYB24 in tomato. Interestingly, homozygous *Slmyb21 Slmyb24* mutant plants showed a drastically impaired flower phenotype, did not produce seeds and retained their petals at the mature fruit (Figure 2D). Ovules of *Slmyb21 Slmyb24* mature flowers showed a similar cell pattern in the innermost cell layer of the nucellus as the *Slmyb21* mutant, characterized by vacuolated cells and accumulation of callose (Supplementary Figure S4, Schubert et al., 2019). Most significantly and distinguished from single *Slmyb21* mutant, flowers of *Slmyb21 Slmyb24* double mutant did not open and showed a severe loss in coloration (Figure 2D). Time series of flower opening/development revealed that the double mutant showed a delayed development and could not open the petals compared to wild type (Supplementary Figure S5, Supplementary Movie 1 and 2). Overall, these results indicate that SlMYB21 and SlMYB24 synergistically regulate flower development beyond female fertility.

**Figure 2.**
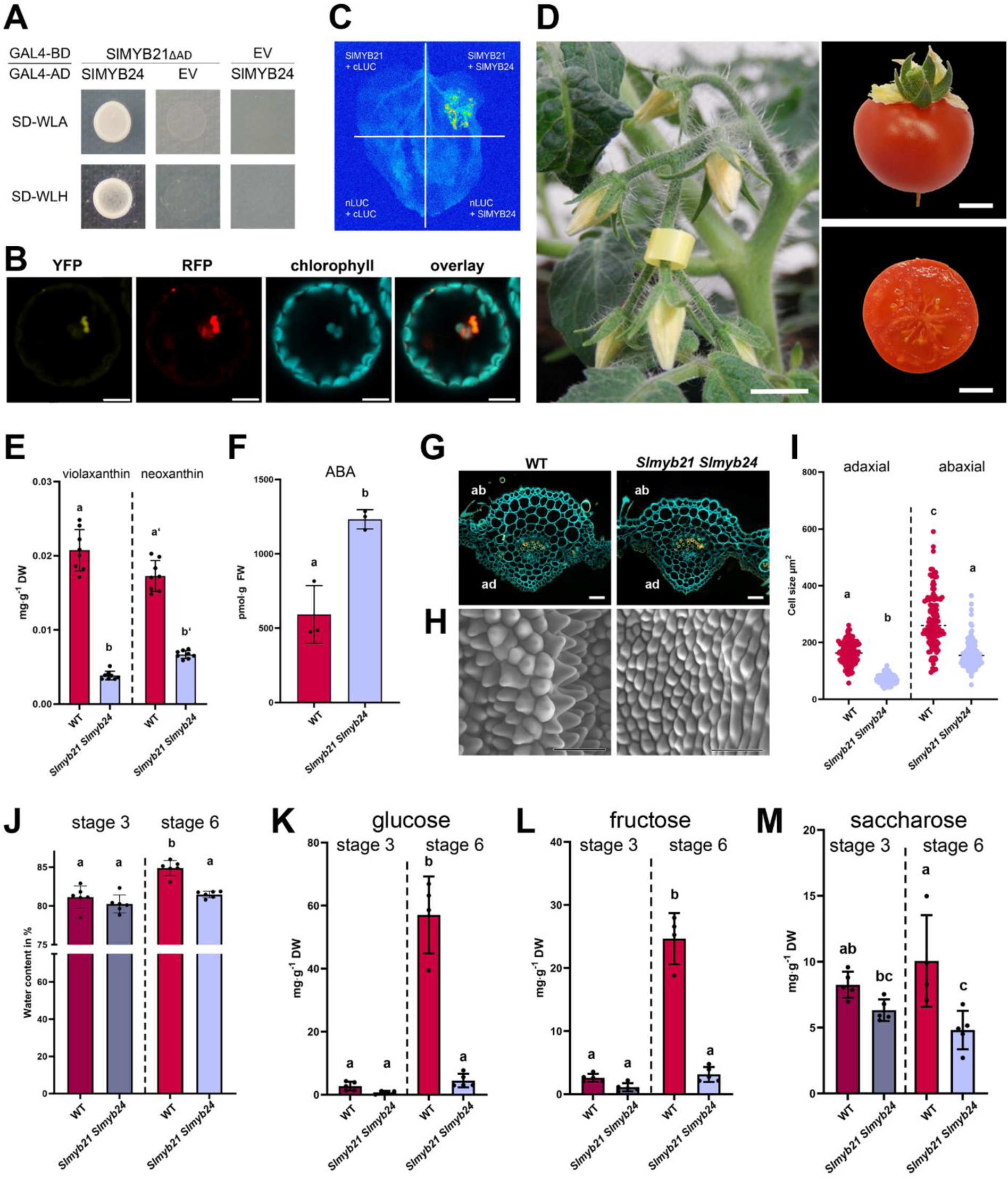
SlMYB21 and SlMYB24 interact and *Slmyb21 Slmyb24* mutant flowers fail to open their petals and do not set seeds. A) Yeast two-hybrid (Y2H) assay assessing SlMYB21-SlMYB24 interaction. The coding sequence of SlMYB21 lacking its activation domain (SlMYB21ΔAD) was fused to the GAL4 DNA-binding domain (BD), and full-length SlMYB24 was fused to the GAL4 activation domain (AD). Yeast was co-transformed with both constructs and maintained on SD/-Trp/-Leu medium. SlMYB21-SlMYB24 interaction was confirmed by growth on SD/-Trp/-Leu/-His and SD/-Trp/-Leu/-Ala. Controls included SlMYB21ΔAD with empty GAL4-BD and empty GAL4-AD with SlMYB24, which did not show growth under selective conditions. B) Bimolecular fluorescence complementation (BiFC) assay confirming SlMYB21-SlMYB24 interaction in *N. benthamiana* protoplasts. *SlMYB21* and *SlMYB24* cDNAs were cloned into 2in1 BiFC vector system to fuse them with N- and C-terminal YFP fragments, respectively. Fluorescence was visualized by confocal laser scanning microscopy 24 h after PEG mediated protoplast transformation. RFP was used as transformation control, chlorophyll autofluorescence is shown in cyan. Scale bars represent 10 µm. C) Split-luciferase complementation assay validating SlMYB21-SlMYB24 interaction. SlMYB21 and SlMYB24 were fused to the C-terminal luciferase fragment (cLuc) and the N-terminal luciferase fragment (nLuc), respectively. Luminescence was recorded 48 h after transient transformation of *N. benthamiana* leaves. Empty vector controls showed no luminescence. Experiments were repeated two times with 3 biological replicates each and showed the same results. D) Floral and fruit phenotypes of homozygous *Slmyb21 Slmyb24* mutants. Mutant flowers exhibit impaired petal expansion/opening, fruits fail to develop viable seeds and still harbor senescent petals. Bars represent 5 mm. E) Petals from *Slmyb21 Slmyb24* flowers exhibit reduced violaxanthin and neoxanthin accumulation at stage 6. Carotenoids were extracted from petals of three flowers per biological replicate. Mean values ± SD (n = 8) are shown. F) Abscisic acid (ABA) levels are elevated in *Slmyb21 Slmyb24* petals at stage 6. ABA was extracted from petals and quantified using LC-MS/MS. Mean values ± SD are shown (n = 3). Different letters in (e-f) indicate significant differences between genotypes as assessed using a two-tailed Student’s *t* test (*P* ≤ 0.05). G) *Slmyb21 Slmyb24* petals exhibit altered morphology of their main vascular bundle at stage 6. Cross-sections were stained with Calcofluor White (CW) and Basic Fuchsin (BF) and visualized by fluorescence microscopy (blue - CW, yellow - BF). ab = abaxial side, ad = adaxial side. Bars represent 50 µm. H) *Slmyb21 Slmyb24* flowers exhibit altered adaxial epidermal cell morphology in petals at stage 6. ESEM micrographs show markedly flattened pavement cells compared to the convex morphology of wild-type petals. Scale bars represent 40 µm. I) Petals from *Slmyb21 Slmyb24* flowers exhibit reduced adaxial and abaxial epidermal cell sizes. Sections from the distal third of petals were prepared, and cell dimensions were quantified from micrographs. Data represent mean ± SD of 120 cells from three independent biological replicates. J) Reduced water content in *Slmyb21 Slmyb24* petals at stage 6. Water content was determined in petals at stage 3 and stage 6. Mean values ± SD are shown (n = 6). K-M) *Slmyb21 Slmyb24* petals exhibit altered carbohydrate levels at stage 6 but not at stage 3. Glucose (K), fructose (L), and sucrose (M) levels were quantified from pools of petals from three flowers per biological replicate. Data represent mean ± SD (*n* ≥ 4). Different letters in (K-M) indicate significant differences between genotypes and developmental stages. Significance was determined by two-way ANOVA followed by Tukey’s HSD post hoc test (*P* ≤ 0.05). Supporting data: Supplemental Figures S4-S6 Supplemental Movies S1 and S2

### SlMYB21 and SlMYB24 regulate petal coloration and opening

Petals from the tomato cultivar MicroTom have a strong yellow color, resulting from the accumulation of carotenoids, mainly violaxanthin and neoxanthin (Galpaz et al., 2006). As the faint yellow color displayed in the flowers of *Slmyb21 Slmyb24* double mutant, we measured the content of violaxanthin and neoxanthin. Both violaxanthin and neoxanthin levels were drastically reduced in *Slmyb21 Slmyb24* petals (Figure 2E). Neoxanthin serves as the precursor of the biosynthesis of the phytohormone abscisic acid (ABA). Therefore, we examined whether ABA accumulation was altered in the petals of the *Slmyb21 Slmyb24* double mutant in comparison to wild type. Surprisingly, we observed an increased accumulation of ABA in petals of *Slmyb21 Slmyb24* (Figure 2F). This contrasts to JA-Ile levels, which were reduced drastically in petals of *Slmyb21 Slmyb24*, pointing again to a positive regulatory function of both TFs on jasmonate biosynthesis (Supplementary Figure S6, Schubert et al., 2019). Thus, these results indicate that SlMYB21 and SlMYB24 promote carotenoid and jasmonate accumulation, while limiting ABA accumulation in petals.

As described above, both *Slmyb21* and *Slmyb21 Slmyb24* flowers exhibited delayed petal opening and showed drastically impaired diurnal movement compared to wild type. Even more notably, *Slmyb21 Slmyb24* flowers displayed closed flowers and completely lost diurnal movement (Supplementary Movie 2). We hypothesized that *Slmyb21 Slmyb24* is arrested at an earlier developmental stage, consequently hindering petal expansion. Histological analysis of petals from developmental stage 6 revealed that the adaxial epidermal cell layer consisted of conical cells in wild type, while the same cells from *Slmyb21 Slmyb24* were flat (Figure 2G, H). Quantification of cell size further showed a reduction in cell size on the adaxial and abaxial sides of *Slmyb21 Slmyb24* petals (Figure 2I). Reversible movement of plant organs is known to be regulated by the movement of water between different tissues (Beauzamy et al., 2014). To test whether the differences in petal movement between wild type and *Slmyb21 Slmyb24* might be caused by differences in water transport, we first measured the total water content of petals. Water content was reduced in *Slmyb21 Slmyb24* petals compared to wild-type petals (Figure 2J). Water pressure, and thus water movement, is mediated by an osmolarity gradient in these tissues (Bieleski et al., 2000). To regulate water gradients, plants utilize osmotically active sugars. In *Slmyb21 Slmyb24* petals, the contents of primary sugars, such as glucose, fructose and sucrose, were reduced in stage 6 compared to wild-type petals (Figure 2K-M). Taken together, these results indicate that SlMYB21 and SlMYB24 synergistically regulate the content of carotenoids and water content as well as soluble sugars in petals.

### Knockout of *SlMYB21* and *SlMYB24* drastically affects transcriptional output

To further elucidate the roles of SlMYB21 and SlMYB24 in petal development and expansion, we conducted transcript profiling using petals collected from wild-type and *Slmyb21 Slmyb24* mutant plants at developmental stages 3 and 6. The transcriptomic data were validated by principal component analysis (PCA), which showed clear separation between wild-type and mutant samples (Supplementary Figure S7). Differential expression analysis was performed by comparing transcriptome of mutant petals with that of wild type at each developmental stage. Using a significance threshold of adj.*p* ≤ 0.05, a total of 3,201 and 10,493 differentially expressed genes (DEGs) were identified at developmental stages 3 and 6, respectively (Supplementary Tables 2 and 3).

To gain insight into the biological functions of these DEGs, Gene Ontology (GO) and MapMan enrichment analyses were performed. At both developmental stages, terms associated with carotenoid biosynthesis and metabolism, including carotene, xanthophyll, and abscisic acid (ABA) biosynthetic processes, were significantly enriched (Figure 3A, Supplementary Tables 4-8). We extracted the DEGs associated with these pathways and examined their expression profiles. In petals of *Slmyb21 Slmyb24*, carotenoid biosynthetic genes, including *PSY2* (Solyc02g081330), *PDS* (Solyc03g123760), *ζ-CAROTENE SATURASE* (*ZDS*, Solyc01g097810), *LCY-B* (Solyc06g074240), *BETA-CAROTENE 3-HYDOXYLASE 2 (BCH2*, Solyc03g007960) were downregulated, whereas several *9-cis-EPOXYCAROTENOID DIOXYGENASEs (NCED)* genes, which catalyze carotenoid cleavage in ABA biosynthesis, were upregulated (Figure 3B, C). The DEGs were also significantly enriched in biological processes associated with petal expansion and movement, including cell wall organization, response to water deprivation, and water transport (Figure 3A, Supplementary Table 4-8).

**Figure 3:**
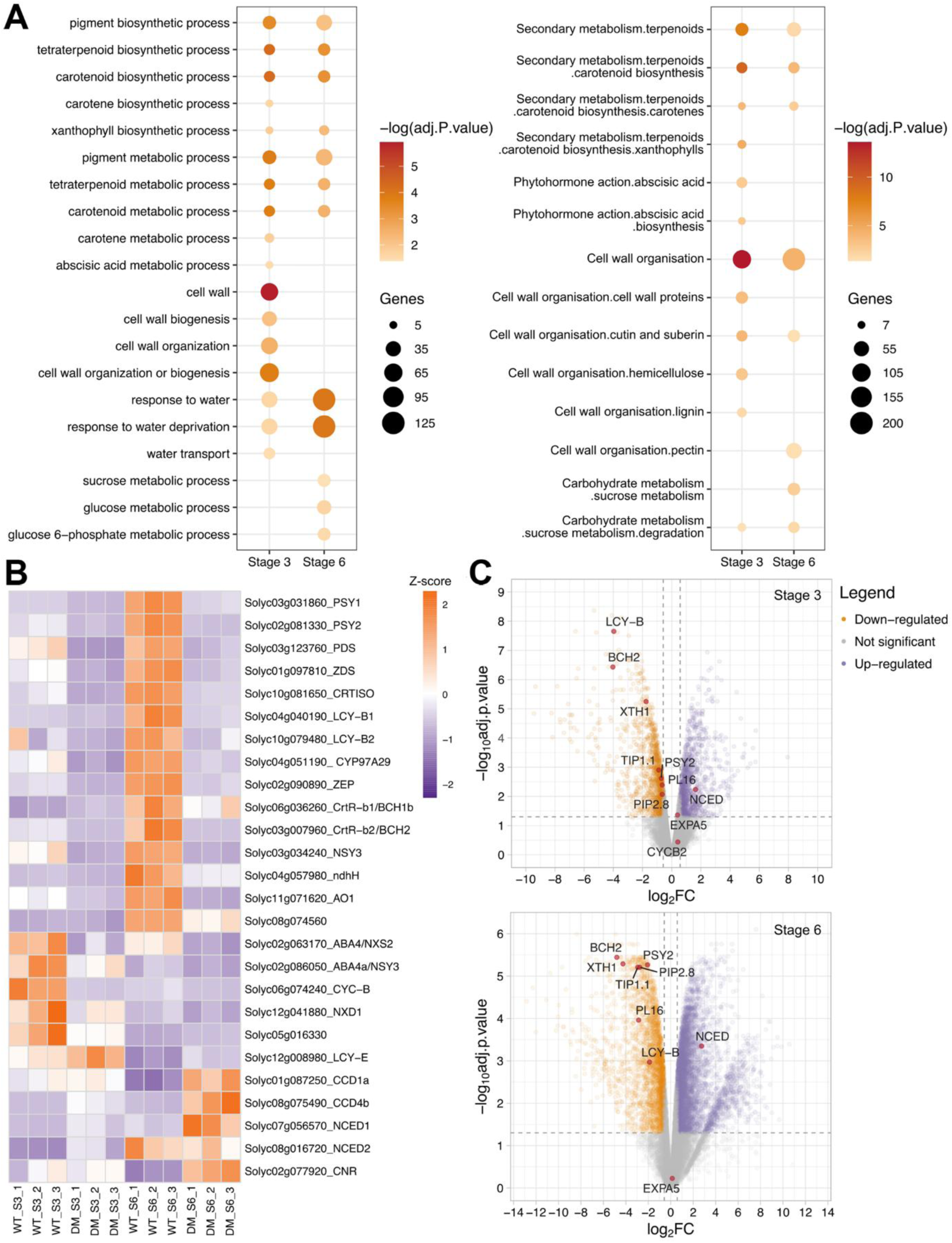
Knockout of *SlMYB21* and *SlMYB24* results in drastic transcriptional changes related to carotenoid biosynthesis and cell wall modifications. A) A selection of significantly enriched terms from Gene Ontology (GO, left panel) and MapMan enrichment (right panel) analyses in differentially expressed genes (DEGs) of *Slmyb21 Slmyb24* double mutants. The -logpval scale represents statistical significance on a continuous color scale from yellow to red. The number of DEGs intersecting with each ontology term is indicated by point size. B) The Transcripts Per Million (TPM) values of carotenoid and abscisic acid (ABA)-related genes intersected in terms associated with carotenoid and ABA biosynthesis and metabolism. The TPM values were retrieved from transcriptomics of petal samples in WT and *Slmyb21 Slmyb24* double mutant (DM) at developmental stage 3 (S3) and 6 (S6), and were presented on a Z-score scale. C) The volcano plots of the transcriptome in *Slmyb21 Slmyb24* petals at developmental stage 3 and 6, compared with WT. The x- and y-axes indicate the log2FC and log10 (adj.P.value), respectively. Yellow and purple dots represent the down-regulated (log2FC < -0.58) and up-regulated genes (log2FC > 0.58), respectively. Grey dots mark non-significantly deregulated genes. The left and right vertical dash lines indicate -0.58 and 0.58. The horizontal dash line delimits the selected log10(0.05) threshold. Supporting data: Supplementary Figure S7

We then confirmed the expression of several carotenoid and ABA-biosynthetic genes. Consistently, expression of the *PSY2* was similar in stage 3 in the *Slmyb21 Slmyb24* mutant as in the wild type but drastically reduced in stage 6 corresponding to the open (wild type) or mature (*Slmyb21 Slmyb24*) flower (Figure 4A). Similarly, the expression of *LCY-B* was drastically reduced in petals of *Slmyb21 Slmyb24* compared to wild type (Figure 4B). Furthermore, transcripts of *BCH2* were almost completely absent in stage 3 and stage 6 of the *Slmyb21 Slmyb24* mutant (Figure 4C). Correspondingly, the transcript levels of *NCED1 (*Solyc07g056570), which encodes the rate-limiting enzyme in ABA synthesis (Thompson et al., 2000) were increased in *Slmyb21 Slmyb24* (Figure 4D). Furthermore, we investigated the expression patterns of genes encoding aquaporins predominantly expressed in petals (Reuscher et al., 2013). The expression of the tonoplast-located aquaporin *TIP1.1* (Solyc06g074820) was not changed at stage 3 but drastically reduced at developmental stage 6 in *Slmyb21 Slmyb24* petals (Figure 4E). The plasma membrane located *PIP2.8* (Solyc01g111660) was similarly downregulated at both developmental stage 3 and 6 in *Slmyb21 Slmyb24* petals (Figure 4F). In addition, enrichment of sucrose- and glucose-related metabolic processes suggests that altered carbohydrate metabolism may further contribute to the defective petal expansion phenotype (Figure 3A, Supplementary Table 4-8). Here, most prominently transcript levels of *LYCOPERSICUM INVERTASE5* (*LIN5,* Solyc09g010080) were drastically reduced in *Slmyb21 Slmyb24* petals in comparison to wild-type petals (Supplementary Figure S8). These results point to a regulation of petal coloration, water transport and sugar metabolism via the transcriptional activity of SlMYB21 and SlMYB24.

**Figure 4.**
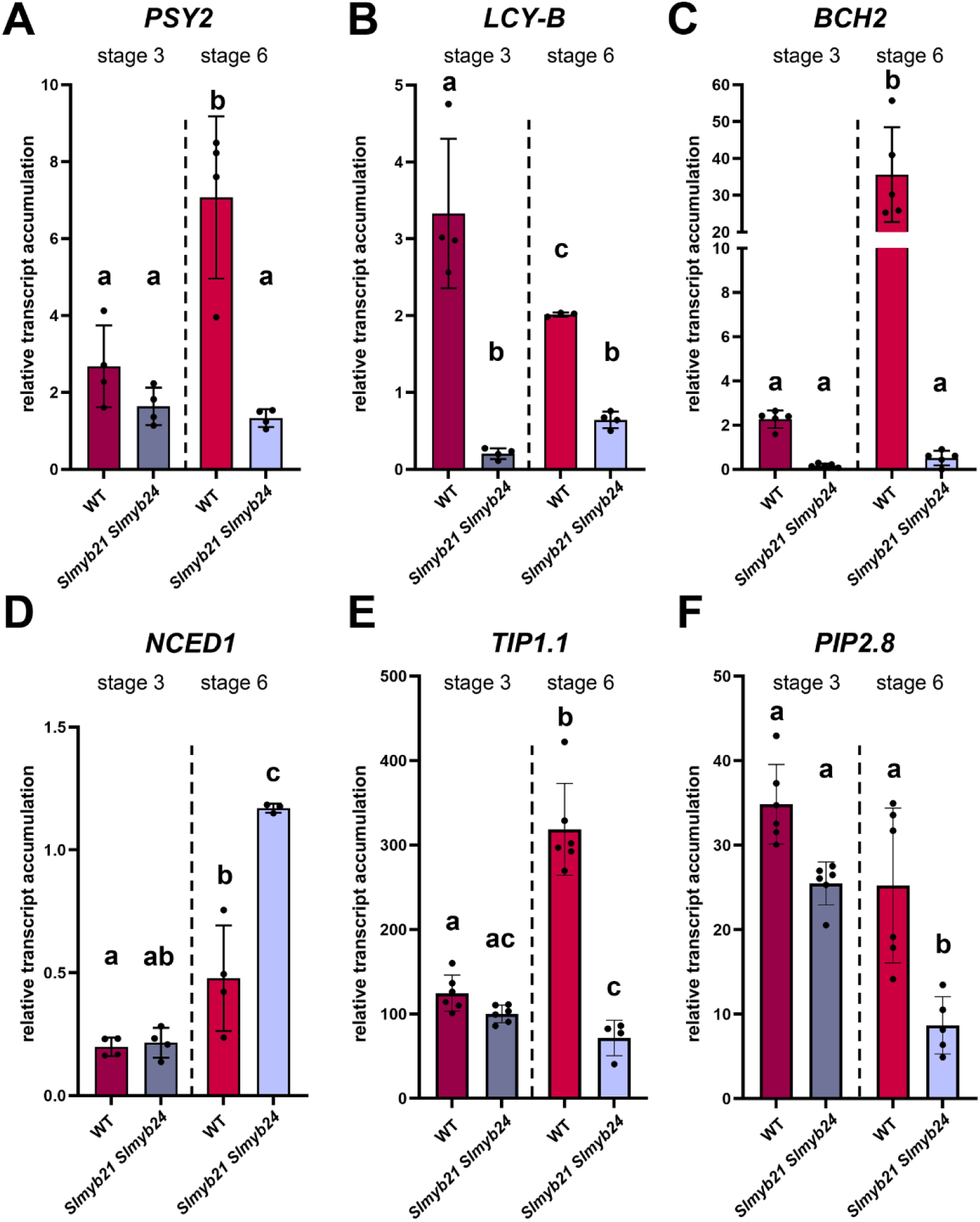
Genes encoding carotenoid biosynthetic enzymes and aquaporins are differentially expressed in *Slmyb21 Slmyb24* mutant petals. Transcript abundance of genes involved in carotenoid biosynthesis, ABA biosynthesis, and water transport: (A) *PSY2*, (B) *LCY-B*, (C) *BCH2*, (D) *NCED1*, (E) *TIP1.1*, and (F) *PIP2.8*. Transcript levels were determined by RT-qPCR and normalized to the reference gene *TIP41*. Data represent means ± SD (*n* ≥ 4). Different letters indicate statistically significant differences among genotypes and developmental stages, as determined by two-way ANOVA followed by Tukey’s honestly significant difference (HSD) post hoc test (*P* ≤ 0.05).

### Genes involved in carotenoid biosynthesis, ABA metabolism, and water transport are regulatory targets of tomato S19-MYBs

To examine the regulatory landscapes of SlMYB21 and SlMYB24 in greater detail, we performed DAP-seq using affinity-purified transcription factors and genomic DNA from tomato cv. MicroTom. Binding peaks for both transcription factors were distributed across upstream, intragenic, and downstream regions of annotated genes (Figure 5A). *De novo* motif analysis of the 600 most significant peaks identified the same core consensus sequence ‘ACCTAAC (reverse complement GTTAGGT),’ for both SlMYB21 and SlMYB24 (Figure 5B). This motif resembles previously described AC-elements (ACC[A/T]A[A/C][T/C] or ACC[A/T][A/C/T][A/C/T]), recognized by S19-MYBs in Arabidopsis (Kelemen et al., 2015), tobacco (LxS-MYB305;) (Liu et al., 2009), and grapevine (VviMYB24; (Zhang et al., 2023). Pearson correlation analysis coefficients further supported the high similarity of DNA-binding motifs among these S19-MYBs (Figure 5C). Notably, approximately 60 % of annotated genes were associated with at least one SlMYB21 or SlMYB24 DAP-seq peak (Figure 5D). Around 55 % of the DAP-seq-associated genes (bound genes) were shared between both TFs, while around 20 % of these shared genes contained binding events within 5kb upstream of their transcriptional start sites (TSS) (Figure 5D). Together, these results reveal highly conserved DNA-binding preferences and extensive overlap in the genomic binding landscapes of SlMYB21 and SlMYB24.

**Figure 5.**
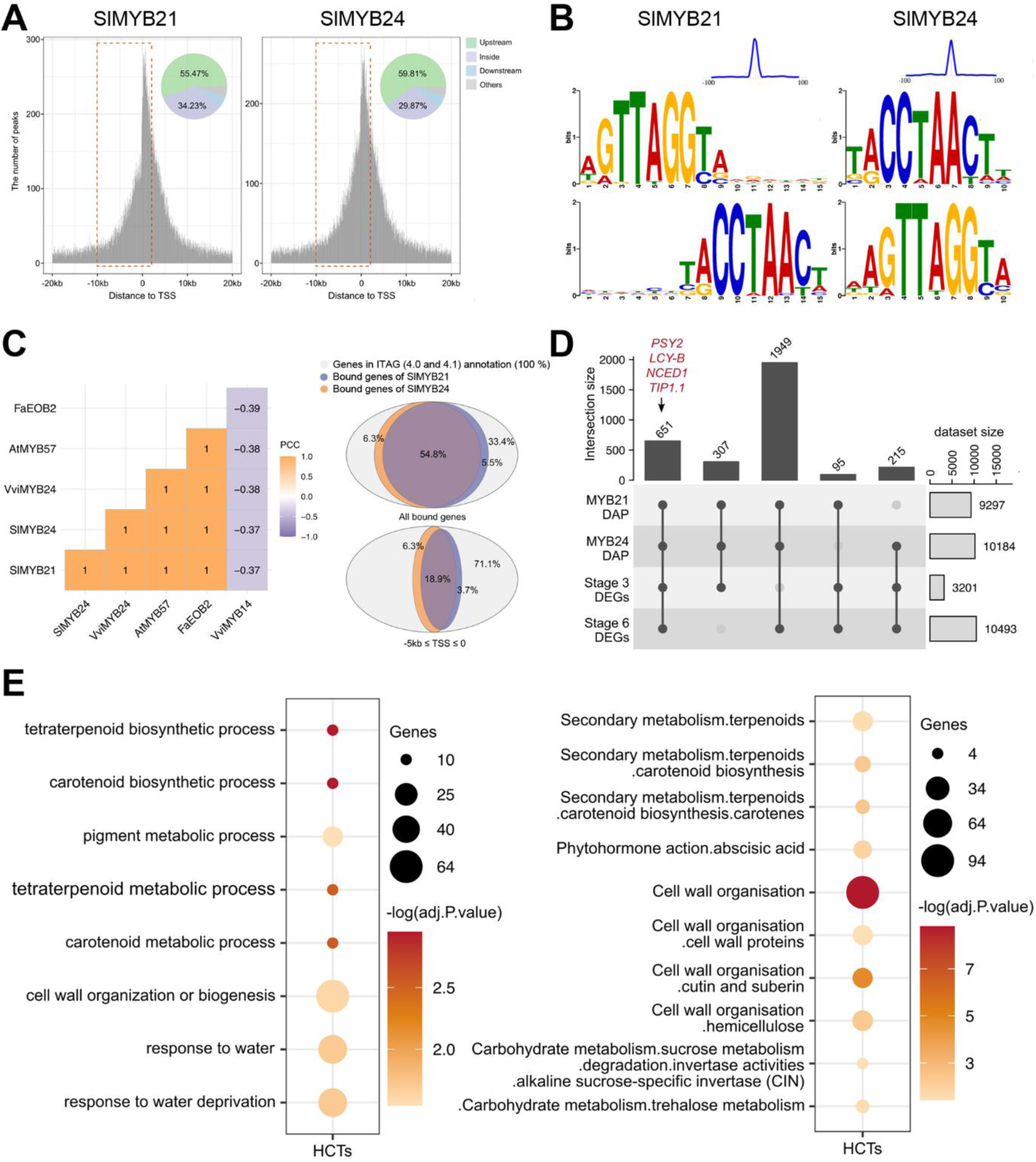
SlMYB21 and SlMYB24 exhibit extensive genome-wide DNA-binding capacity and share a substantial proportion of their associated genes. A) Distribution of SlMYB21 (left panel) and SlMYB24 (right panel) DAP-seq binding peaks relative to the transcription start sites (TSSs) of annotated genes in the tomato cv. Heinz reference genome. B) DNA-binding motifs identified from the 600 highest-ranking DAP-seq peaks for SlMYB21 (left panel) and SlMYB24 (right panel), shown in forward (top) and reverse-complement (bottom) orientations. The blue curve in the upper-right panel represents the positional enrichment of the identified motif within the sequences corresponding to the 600 highest-ranking peaks. C) Comparison of consensus DNA-binding motifs identified from the top 600 highest-ranking peaks of subgroup 19 (S19) R2R3-MYB transcription factors from *Arabidopsis thaliana*, *Solanum lycopersicum*, *Fragaria* x *ananassa*, and *Vitis vinifera.* The consensus motif identified for the R2R3-MYB transcription factor VviMYB14 (Orduña et al., 2022), was included as a reference outside the S19 subgroup. Motif similarity was assessed using Pearson correlation coefficients (PCCs) and is represented by a color gradient from purple to yellow. D) Comparison of the genome-wide binding repertoires of SlMYB21 and SlMYB24, illustrating the overlap between their DAP-seq-associated genes in the tomato cv. Heinz reference genome and the differentially expressed genes (DEGs) identified in *Slmyb21 Slmyb24* double-mutant flowers at developmental stages 3 and 6 relative to wild type (WT). E) Selected significantly enriched Gene Ontology (GO) terms associated with genes identified in the SlMYB21 (left panel) and SlMYB24 (right panel) DAP-seq datasets. Dot size represents the number of genes associated with each GO term, while color indicates enrichment significance, expressed as −log₁₀(*P*-value), on a continuous yellow-to-red scale.

To identify candidate regulatory targets of SlMYB21 and SlMYB24, we integrated their DAP-seq binding profiles with transcriptomic data from *Slmyb21 Slmyb24* petals. For this analysis, we focused on promoter-associated binding events, operationally defined as peaks located between 5 kb upstream and 200 bp downstream of the TSS. This criterion was selected to prioritize genes with promoter-associated binding events that could be investigated using conventional promoter-based reporter assays. The additional 200 bp downstream of the TSS were included to retain binding peaks spanning the promoter–5′ UTR boundary, which might otherwise be excluded by a strictly upstream cutoff. Accordingly, the resulting high-confidence targets (HCTs) represent a promoter-associated subset of the broader SlMYB21 and SlMYB24 binding repertoire, which also includes peaks within gene bodies and downstream regions.

Using this promoter-proximal criterion, we identified 9,297 and 10,184 DAP-seq-associated genes for SlMYB21 and SlMYB24, respectively (Supplementary Tables 9 and 10). These datasets were integrated with the 3,201 and 10,493 DEGs (adjusted P ≤ 0.05) identified in double-mutant petals at developmental stages 3 (S3) and 6 (S6), respectively (Supplementary Tables 2 and 3). This yielded 3,217 high-confidence promoter-associated targets supported by at least three of the four datasets. Among these promoter-associated HCTs, 651 genes were represented in all four datasets, constituting the most stringent shared target set (Figure 6A; Supplementary Table 11). GO and MapMan enrichment analyses of the 3,217 HCTs revealed significant enrichment of functional categories associated with carotenoid biosynthesis and metabolism, cell wall organization, responses to water deprivation, and sucrose metabolism (Figure 6B; Supplementary Tables 12 and 13).

**Figure 6:**
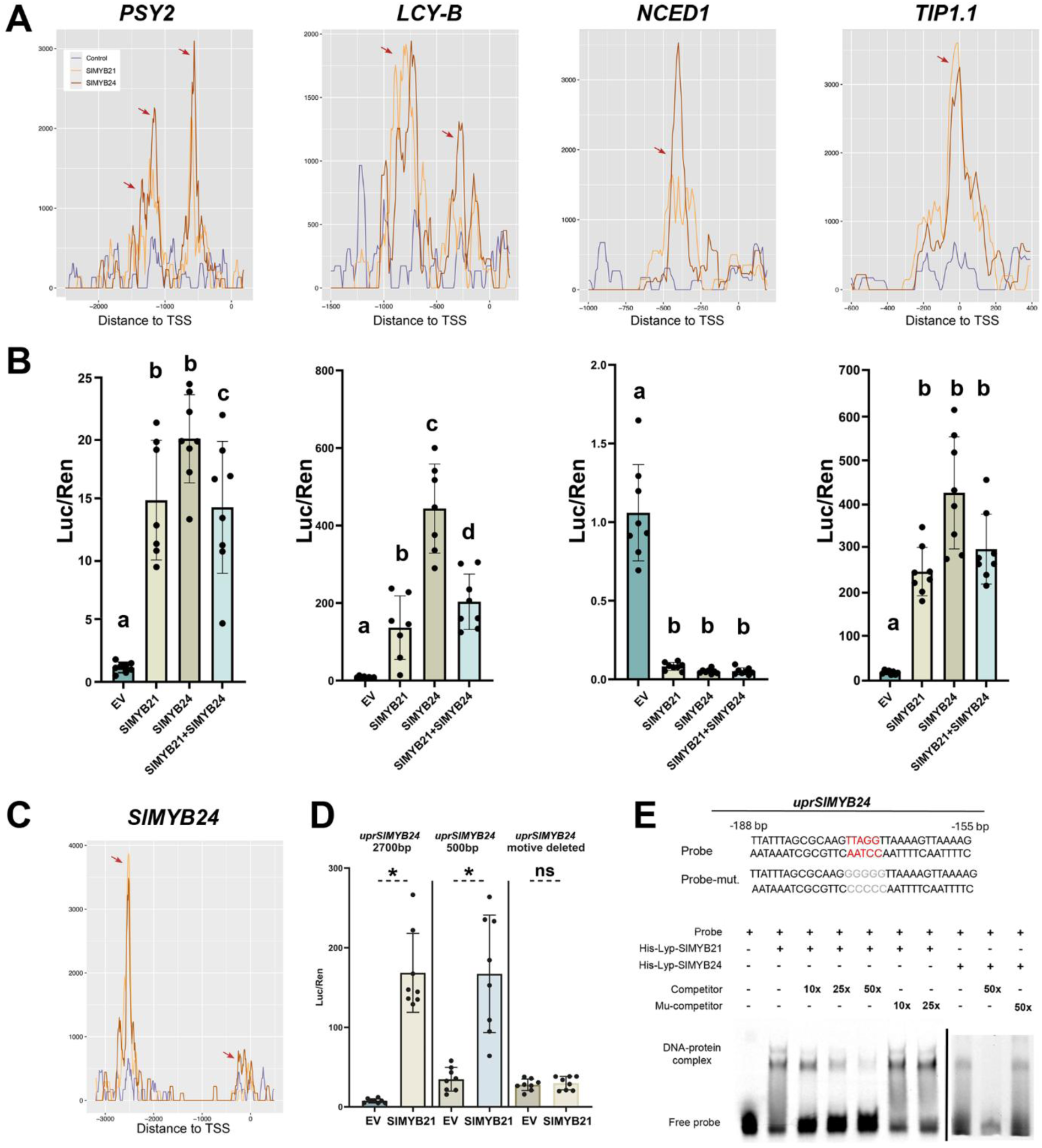
SlMYB21 and SlMYB24 bind and regulate the promoters of genes involved in carotenoid biosynthesis, ABA homeostasis, and water transport. A) DAP-seq binding profiles of SlMYB21 and SlMYB24 in the upstream regulatory regions (*upr*) of *PSY2*, *LCY-B*, *NCED1*, and *TIP1.1*, showing the positions of the identified binding peaks. B) Transient dual-luciferase reporter assays assessing the effects of SlMYB21 and SlMYB24, individually or in combination, on the activity of the selected upstream regulatory regions. Promoter activity is expressed as the fold change in the firefly luciferase (LUC)/Renilla luciferase (REN) luminescence ratio relative to the empty-vector (EV) control. C) DAP-seq binding profile of SlMYB21 at the *SlMYB24* upstream regulatory region (*uprSlMYB24*), showing two identified binding peaks. D) Transient dual-luciferase reporter assays using truncated and mutated versions of *uprSlMYB24* to assess SlMYB21-dependent transcriptional activation. Promoter activity is expressed as the fold change in the LUC/REN luminescence ratio relative to the EV control. A 500-bp upstream fragment was sufficient to support SlMYB21-dependent activation, whereas mutation of the AC-rich binding element abolished this response. In (B) and (D), values represent means ± SD (*n* ≥ 6). Different letters indicate statistically significant differences according to two-way ANOVA followed by Tukey’s honestly significant difference (HSD) post hoc test (*P* ≤ 0.05). E) Electrophoretic mobility shift assay (EMSA) showing binding of recombinant His-Lyp-SlMYB21 and His-Lyp-SlMYB24 fusion proteins to a *uprSlMYB24* DNA fragment containing the GTTAGGT motif. The mutated probe (probe mut.) carries substitutions within the TTAGG core sequence, indicated in red. Competition assays were performed using unlabeled wild-type probe at 10-, 25-, and 50-fold molar excess or unlabeled mutated probe (mut. competitor) at 10- and 25-fold molar excess relative to the labeled probe. Supporting data: Supporting Figures S8-S11

Among the 651 genes supported by all four datasets, *PSY2*, *LCY-B*, *NCED1*, and *TIP1.1* emerged as candidate direct regulatory targets of SlMYB21 and SlMYB24. All four genes displayed DAP-seq binding peaks for both transcription factors within 2 kb upstream of their TSSs (Figure 6A; Supplementary Table 11). To investigate their transcriptional regulation, we performed transient promoter activation assays in *N. benthamiana* leaves. Consistent with the transcriptomic data, both SlMYB21 and SlMYB24 strongly activated the *uprs* of *PSY2*, *LCY-B*, and *TIP1.1*, whereas *NCED1 upr* activity was significantly repressed (Figure 6B). Interestingly, although no clear DAP-seq binding peaks were detected in the *LIN5 upr*, both transcription factors strongly enhanced its activity in transient assays (Supplementary Figure S9), suggesting that *LIN5* may be regulated indirectly or through mechanisms not captured by the DAP-seq analysis. Together, these findings support a role for SlMYB21 and SlMYB24 in positively regulating genes involved in carotenoid biosynthesis and water transport, while negatively regulating *NCED1*, a key gene involved in ABA biosynthesis.

### SlMYB21 directly activates *SlMYB24* transcription

Expression profiling across tomato tissues and organs revealed that *SlMYB21* and *SlMYB24* exhibit similar flower-enriched expression patterns. However, *SlMYB24* transcript accumulation occurred later than that of *SlMYB21* during flower development (Supplementary Figure S10). Furthermore, *SlMYB24* expression was nearly abolished in *Slmyb21* mutant carpels, suggesting that SlMYB21 may act upstream of *SlMYB24* in the transcriptional regulatory network controlling flower development (Supplementary Figure S11).

To investigate this potential regulatory relationship, we first examined the ability of SlMYB21 to bind to the *upr* of *SlMYB24*. DAP-seq analysis identified two SlMYB21 binding peaks located 2,564 and 174 bp upstream of the *SlMYB24* TSS, respectively (Figure 6C). We next evaluated the transcriptional activity of SlMYB21 using transient promoter activation assays, which revealed a significant increase in *SlMYB24 upr* activity in the presence of SlMYB21 (Figure 6D).

To determine whether this activation depends on the conserved AC-element recognized by SlMYB21, we introduced mutations into the GTTAGGT motif located within the 500-bp *SlMYB24 upr* fragment immediately upstream of the TSS. Disruption of this motif abolished SlMYB21-dependent *upr* activation, reducing luciferase activity to levels similar to those of the empty-vector control (Figure 6D). Electrophoretic mobility shift assays (EMSAs) further confirmed direct binding of SlMYB21 and SlMYB24 to the identified AC-element (Figure 6E). Together, these results demonstrate that SlMYB21 recognizes an AC-rich *cis*-regulatory element in the *SlMYB24* promoter and positively regulates its transcription, establishing a direct hierarchical regulatory relationship between these two S19-MYB transcription factors.

## Discussion

Flower opening and movement is a complex developmental process coordinated by endogenous developmental cues, circadian rhythm, and phytohormonal signaling. Several studies have shown that flower development and fertility are tightly controlled by jasmonates, as demonstrated by the sterility phenotypes of JA-deficient and JA-insensitive mutants. Interestingly, the reproductive consequences of impaired JA signaling differ between species: whereas the corresponding *A. thaliana* mutants exhibit male sterility, orthologous tomato mutants display female sterility (Wasternack and Hause, 2013). Beyond fertility, JA signaling also contributes to petal development, as evidenced by impaired flower opening in JA-related mutants. Two important transcriptional regulators mediating JA-dependent floral development are MYB21 and MYB24, whose functions have been investigated in both Arabidopsis and tomato. Here, we demonstrate that the tomato homologs SlMYB21 and SlMYB24 have overlapping but unequal contributions to petal pigmentation and opening. Whereas *Slmyb24* flowers are largely indistinguishable from wild-type flowers, *Slmyb21* mutants exhibit pale petals and pronounced defects in flower opening, which are further exacerbated in the *Slmyb21 Slmyb24* double mutant. This unequivocally proves that the role of both transcription factors is not restricted to the regulation of fertility in the ovaries of tomato flowers, as originally demonstrated for SlMYB21 (Schubert et al., 2019), but extends to the coordinated regulation of petal development, pigmentation, and movement.

Petal pigmentation provides important visual cues for pollinator attraction, and flowering plants have evolved a remarkable diversity of pigmentation patterns. In tomato, the characteristic yellow coloration of petals is primarily determined by carotenoid accumulation, particularly xanthophylls, such as neoxanthin and violaxanthin (Isaacson et al., 2002). Beyond pigmentation, carotenoids also contribute to photoprotection. They are known, especially neoxanthin, to scavenge singlet oxygen formed within the chloroplast under highlight stress conditions (Young, 1991). We observed a pronounced reduction in neoxanthin and violaxanthin accumulation in *Slmyb21 Slmyb24* petals, accompanied by reduced expression of several carotenoid biosynthetic genes. Carotenoid metabolism in tomato is governed by a network of enzymes that display pronounced tissue–specific expression patterns. For example, the *BCH2* gene is highly expressed in petals and its disruption results in a white–flower phenotype (Galpaz et al., 2006). Although *BCH2* expression was strongly reduced in the double mutant, our DAP-seq analysis did not identify promoter-associated binding by SlMYB21 and SlMYB24, suggesting that its transcriptional downregulation may be indirect. In contrast, we identified *PSY2* and *LCY-B/NSY* as candidate direct targets of both transcription factors, supported by promoter-associated DAP-seq peaks and transcriptional activation in transient reporter assays (Figure 6). These findings are consistent with previous reports of preferential *PSY2* and *LCY-B* expression in floral tissues (Ronen et al., 2000; Giorio et al., 2008) and indicate that SlMYB21 and SlMYB24 positively regulate carotenoid accumulation through multiple steps of the biosynthetic pathway. Nevertheless, their contribution is unlikely to be exclusive, as additional transcriptional regulators and indirect effects may participate in establishing the petal carotenoid profile. Also, it remains elusive why these TFs regulate the expression of enzymes at different steps in carotenoid-biosynthesis. An intriguing possibility is that SlMYB21- and SlMYB24-dependent carotenoid accumulation contributes not only to visual attraction but also to the protection of reproductive tissues against excessive irradiance. Such a dual role would be consistent with the previously described involvement of grapevine VviMYB24 in the regulation of terpenes and flavonols in grape berries under high light stress conditions (Zhang et al., 2023), suggesting that S19-MYBs may have retained broader functions linking specialized metabolism with environmental responses.

In addition to promoting carotenoid accumulation, SlMYB21 and SlMYB24 seem to contribute to ABA homoeostasis in developing petals. Both TFs reduced *NCED1* promoter activity in transient assays, consistent with the elevated ABA levels observed in *Slmyb21 Slmyb24* petals. Because NCED enzymes catalyze a key step in ABA biosynthesis, these observations support a model in which SlMYB21 and SlMYB24 negatively regulate ABA accumulation, at least partly through transcriptional repression of *NCED1*. The most prominent mechanism regulated by ABA is a central regulator of plant water relations and is also involved in developmental processes, including organ senescence and abscission. This is in line with our findings that *Slmyb21 Slmyb24* petals exhibited reduced water content and failed to undergo normal abscission during fruit development (Figure 2). Although these phenotypes cannot be attributed exclusively to increased ABA, they suggest a possible connection between altered hormonal homeostasis and the physiological defects observed in the mutant.

Moreover, ABA has been reported to exert a modest inhibitory effect on flower opening in Iris (van Doorn et al., 2013). Therefore, elevated ABA accumulation may contribute to impaired petal opening in *Slmyb21 Slmyb24*, potentially acting alongside alterations in water transport and osmotic regulation.

In addition to pigmentation, flower opening is important for reproduction, often through the presentation of petals to attract animals that promote pollination (van Doorn and Kamdee, 2014). Flower opening requires coordinated changes in petal growth, cellular water relations, and tissue mechanics. Petal development generally involves an initial phase dominated by cell proliferation, followed by cell expansion (Krizek and Anderson, 2013). Whereas petal growth involves the expression of genes associated with cell-wall modification (Ochiai et al., 2013), the final opening process can depend on changes in cellular osmotic pressure and water uptake in cells that do not elongate, leading to differential expansion and movement of petal tissues (van Doorn and Kamdee, 2014). These processes are influenced by soluble sugar availability and aquaporin activity, which together contribute to the regulation of cellular water potential and transmembrane water transport (Abascal et al., 2014). Aquaporins function is regulated at multiple levels, including post-translational modifications and diurnal phosphorylation cycles (Nemoto et al., 2022). Nevertheless, transcriptional regulation may also influence the capacity of petal cells to redistribute water during opening. In this context, the decreased expression of aquaporin-encoding genes *TIP1.1* and *PIP2.8* in *Slmyb21 Slmyb24* petals is particularly relevant. We identified both aquaporins as a promoter-associated targets of both SlMYB21 and SlMYB24, all of which show their highest expression in fully opened petals. Transient assays demonstrated activation of their *uprs* by both factors, supporting a direct regulatory connection between these S19-MYBs and cellular water transport.

As TIP1.1 is predicted to localize to the tonoplast, its reduced expression could impair vacuolar water transport and turgor changes required for petal movement. We therefore propose that SlMYB21- and SlMYB24-dependent activation of *TIP1.1* contributes to water redistribution during flower opening, potentially influencing differential expansion between petal surfaces (Figure 7). This mechanism remains hypothetical and will require direct measurements of cellular water dynamics and aquaporin activity. In turn, the reduced water content of double-mutant petals was also accompanied by decreased soluble sugar levels, suggesting that impaired osmotic regulation may contribute to the opening defect. In particular, reduced expression of *LIN5*, which encodes a cell-wall invertase involved in phloem unloading and sink metabolism, could contribute to altered sugar availability (Godt and Roitsch, 1997; Zanor et al., 2009). These findings are consistent with recent findings in *Nicotiana attenuata,* where corolla movement is driven by water redistribution movement and cell expansion, rather than increased cell proliferation (Choi et al., 2026).

**Figure 7.**
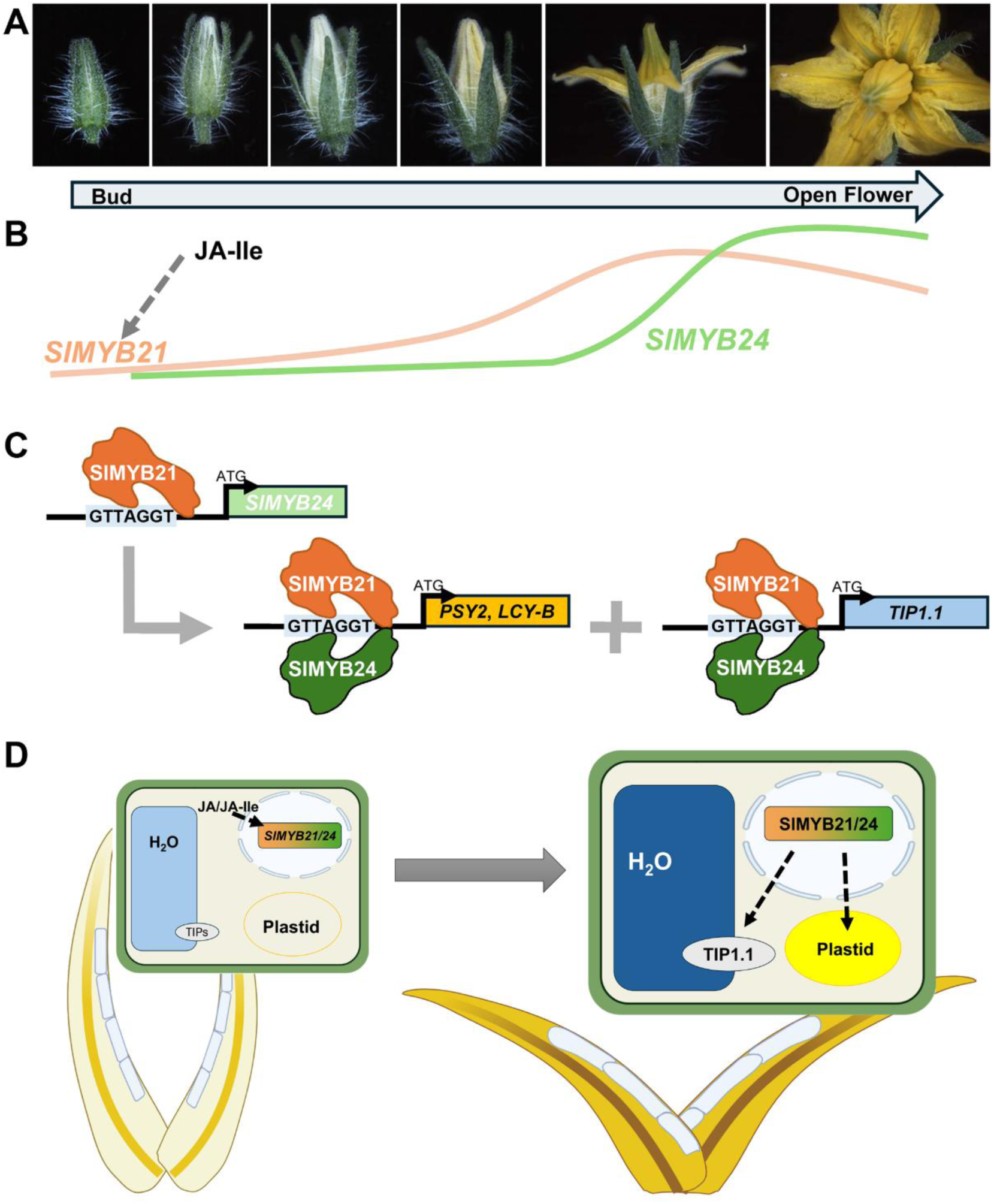
Model summarizing the role of SlMYB21 and SlMYB24 in petal development of tomato. A) Representative images of flower buds and open flowers from wild-type tomato cv. Micro-Tom illustrating the six developmental stages. B) Schematic representation of the proposed temporal relationship between endogenous JA-Ile accumulation and the expression of *SlMYB21* and *SlMYB24* during flower development. Increased JA-Ile levels are associated with the induction of *SlMYB21*, followed by the accumulation of *SlMYB24* transcripts. C) Proposed transcriptional regulatory network involving SlMYB21 and SlMYB24. SlMYB21 recognizes the AC-rich cis-regulatory element (GTTAGGT/ACCTAAC) in the upstream regulatory region of *SlMYB24* and activates its transcription. Both SlMYB21 and SlMYB24 can recognize similar AC-rich elements and regulate overlapping sets of downstream target genes, including genes involved in carotenoid biosynthesis, ABA homeostasis, and water transport. The two TFs may bind these elements independently and/or as part of a protein complex. D) Proposed model linking SlMYB21- and SlMYB24-dependent transcriptional regulation to petal maturation. During early flower development, when *SlMYB21* and *SlMYB24* expression is low, petals remain pale and closed. As development progresses, increased JA/JA-Ile signaling promotes the expression of these TFs, which in turn regulate genes involved in carotenoid accumulation, ABA homeostasis, and cellular water transport. These coordinated processes contribute to yellow pigmentation, changes in cellular water content and turgor, and ultimately petal expansion and flower opening. This figure was created with BioRender (www.biorender.com).

A further contribution of this study is the genome-wide identification of the ‘GTTAGGT’ (‘ACCTAAC’) consensus motif as the most frequent DNA-binding site of SlMYB21 and SlMYB24 in the tomato genome (Figure 5). SlMYB21 and SlMYB24 belong to the R2R3-MYB family TFs which constitute one of the largest TF gene families in plant genomes, e.g., with approximately 127 members reported in tomato (Millard et al., 2019; Liu et al., 2026). R2R3-MYB proteins regulate diverse biological processes, including plant specialized metabolism, environmental responses and plant growth and development (Wu et al., 2022). Their involvement in reproductive development extends across flowering plants, with reported function in petal and pistil development in *Petunia axillaris* (Chopy et al., 2023), inflorescence architecture and the number of flowers in chickpea (*Cicer arietinum*) (Caballo et al., 2022), and anther development and pollen formation in *Lilium* species (Liu et al., 2021). Previous studies have shown that R2R3-MYB proteins recognize several *cis*-regulatory elements, including MYB-core motifs [C/T]NGTT[G/T] and AC-rich elements (Prouse and Campbell, 2012; Franco-Zorrilla et al., 2014; Kelemen et al., 2015). Our identification of a conserved AC-rich motif for SlMYB21 and SlMYB24 indicates that these closely related S19-MYBs retain highly similar intrinsic DNA-binding preferences. Their extensive overlap in DAP-seq-associated genes further supports this interpretation. Importantly, our DAP-seq analyses revealed that SlMYB21 and SlMYB24 binding events were not restricted to *uprs* but were also distributed within gene bodies and downstream of annotated genes. Although we focused on a −5 kb/+200 bp window relative to the TSS to define promoter-associated HCTs and prioritize candidates for functional validation, this operational criterion does not necessarily encompass the full regulatory repertoire of either transcription factor. Indeed, functional *cis*-regulatory elements can also occur within introns, other transcribed regions, and downstream sequences, potentially contributing to transcriptional regulation through mechanisms that are not captured by conventional promoter-based assays. Consequently, genes associated with binding events outside our selected window may represent additional regulatory targets, particularly when supported by differential expression in the mutant. Nevertheless, DAP-seq identifies potential DNA-binding sites *in vitro*, and additional evidence would be required to establish whether these distal or intragenic interactions contribute to transcriptional regulation *in planta*. Future integration of chromatin accessibility and chromatin conformation data, together with targeted functional assays, could help resolve the regulatory significance of these binding events.

Despite their similar DNA-binding preferences and extensive overlap in candidate targets, SlMYB21 and SlMYB24 make markedly different contributions to tomato flower development. Whereas loss of SlMYB24 alone produces no obvious floral phenotype, disruption of SlMYB21 causes pronounced defects that become more severe in the double mutant. Our finding that SlMYB21 directly binds to and activates the *SlMYB24* promoter provides a mechanistic explanation for at least part of this functional asymmetry. Loss of SlMYB21 may compromise both its own regulatory activity and the expression of SlMYB24, whereas loss of SlMYB24 leaves SlMYB21 function intact, potentially allowing compensation through overlapping target regulation. The enhanced phenotype of the double mutant nevertheless indicates that SlMYB24 retains a biologically relevant contribution that becomes apparent in the absence of SlMYB21. At the same time, the possibility that SlMYB21 controls additional targets independently of SlMYB24 cannot be excluded. Thus, the relationship between these transcription factors appears to involve both hierarchical transcriptional regulation and partially overlapping downstream functions, providing a plausible explanation for their unequal genetic contributions despite their conserved DNA-binding specificity.

Our findings collectively support a regulatory model in which SlMYB21 and SlMYB24 coordinate carotenoid biosynthesis and water transport to control petal pigmentation and opening of tomato flowers (Figure 7). The proposed model also distinguishes SlMYB21 positively regulating *SlMYB24* transcription, while both proteins can recognize similar AC-rich *cis*-regulatory elements and regulate overlapping sets of downstream genes. Importantly, their physical interaction does not necessarily imply that SlMYB24 requires SlMYB21 for DNA recruitment, as DAP-seq demonstrates that SlMYB24 can independently recognize the AC-element *in vitro*. The two transcription factors may therefore associate with shared regulatory sequences independently and/or as components of a protein complex, possibilities that remain to be resolved experimentally. Nevertheless, our data offers new insights into the role of these two R2R3-MYB TFs within the transcriptional regulatory network governing tomato floral shape and color and provides a mechanistic framework for understanding how closely related regulators coordinate distinct physiological processes during floral development.

## Acknowledgment

We thank Claus Wasternack (IPB Halle) for critical reading the manuscript; Hagen Stellmach (IPB Halle) for assistance with hormone measurements and embeddings; Thomas Vogt and Osmany Jose Diaz Bravo (IPB Halle) for their support with carotenoid measurements; Simone Fraas (Martin-Luther-University Halle-Wittenberg) for sectioning carpels; and Konrad Koller (IPB Halle) for assistance with video recordings. We are grateful to Prof. Z. Shakked (Weizmann Institute) for providing the pHLT plasmid. We also thank Pablo Romero (I2SysBio) for his assistance with greenhouse plant maintenance and *in vitro* plant regeneration.

## Author contributions

M.F., C.Z., J.T.M. and B.H. conceptualized and designed the research. H.S. and C.Z. identified the *SlMYB24* sequence. H.S. generated the *Slmyb24* and *Slmyb21 Slmyb24* mutants. M.F., C.Z., H.S., J.D.H., L.H., I.B., S.W. and J.E. performed the experiments. A.S. analyzed data. F.S. took the ESEM pictures. M.F., C.Z. and B.H. wrote the first draft of the manuscript. B.H. and J.T.M. supervised the research and acquired funding. All authors have reviewed and approved this submission.

## Supplementary Material

Supplementary Figure S1: SlMYB24 is the ortholog of AtMYB24.

Supplementary Figure S2: Sequence alignment of *Slmyb24-3* to genomic SlMYB24 sequence unveil a premature stop codon.

Supplementary Figure S3: *SlMYB21* expression shows only minor changes in carpels and petals of *Slmyb24* in comparison to wild type.

Supplementary Figure S4: Ovules of *Slmyb21 Slmyb24* at stage 6 show altered morphology and PCD in the nucellus.

Supplementary Figure S5: *Slmyb21, Slmyb21 Slmyb24* but not *Slmyb24* exhibit delayed flower development.

Supplementary Figure S6: Jasmonate levels in petals.

Supplementary Figure S7: Principal component analysis (PCA) of transcriptomic data from petals of WT and *Slmyb21 Slmyb24* double mutants at developmental stage 3 and 6.

Supplementary Figure S8: *LIN5* transcript levels are impaired in petals from *Slmyb21 Slmyb24*.

Supplementary Figure S9: *uprLIN5* is activated by SlMYB21 and SlMYB24.

Supplementary Figure S10: *SlMYB24* expression follows the *SlMYB21* expression during petal development.

Supplementary Figure S11: *SlMYB24* expression is almost not detectable in the *Slmyb21* mutant.

Supplementary data 1: Sequences of the *upr* regions of selected tomato genes. Supplementary Table S1: List of primers used.

Supplementary Table S2: Differentially expressed genes at developmental stage 3. Supplementary Table S3: Differentially expressed genes at developmental stage 6.

Supplementary Table S4: Gene ontology analysis of differentially expressed genes at stage 3.

Supplementary Table S5: Gene ontology analysis of differentially expressed genes at stage 6.

Supplementary Table S6. MapMan enrichment analysis of differentially expressed genes at stage 3.

Supplementary Table S7: MapMan enrichment analysis of differentially expressed genes at stage 6.

Supplementary Table S8: TPM values generated by transcriptome of petal samples collected from WT and *Slmyb21 Slmyb24* double mutants (DM) at developmental stage 3 and 6.

Supplementary Table S9: SlMYB21-DAP results mapped in Heinz ‘SL4.0’ reference genome assembly, filtered by peak position (−5kb ≤ distance to TSS ≤ 0.2kb).

Supplementary Table S10: SlMYB24-DAP results mapped in Heinz ‘SL4.0’ reference genome assembly, filtered by peak position (−5kb ≤ distance to TSS ≤ 0.2kb).

Supplementary Table S11: High-Confidence Targets (HCTs) of S19-SlMYBs identified by integrating DAP-seq and RNA-seq: genes bound by SlMYB21 and SlMYB24, and differentially expressed genes (DEGs) in *Slmyb21 Slmyb24* petals at developmental stages 3 (S3) and 6 (S6).

Supplementary Table S12: Gene ontology analysis of S19-SlMYBs High-Confidence Targets.

Supplementary Table S13: MapMan enrichment analysis of S19-SlMYBs High-Confidence Targets.

Supplemental Movie 1: Petal movement of WT flowers follows the diurnal rhythm. Supplemental Movie 2: In *Slmyb21 Slmyb24* flowers petal movement is absent.

## Data availability statement

The transcriptomic raw data and DAP-seq data generated in this study have been deposited in Sequence Read Archive (SRA) of NCBI under the accession numbers PRJNA1537550 and PRJNA1539859, respectively.

