## Supplemental figures for "SlMYB21 and SlMYB24 cooperatively regulate petal coloration and movement in *Solanum lycopersicum*"

to

by

Moritz Friesch, Chen Zhang, Henrikje Smits, Julius Decker-Hibbel,
Lena Harborth, Saskia Weinmar, Isabel Bühler, Antonio Santiago,
Jone Echeverría, Frank Syrowatka, José Tomás Matus, Bettina
Hause

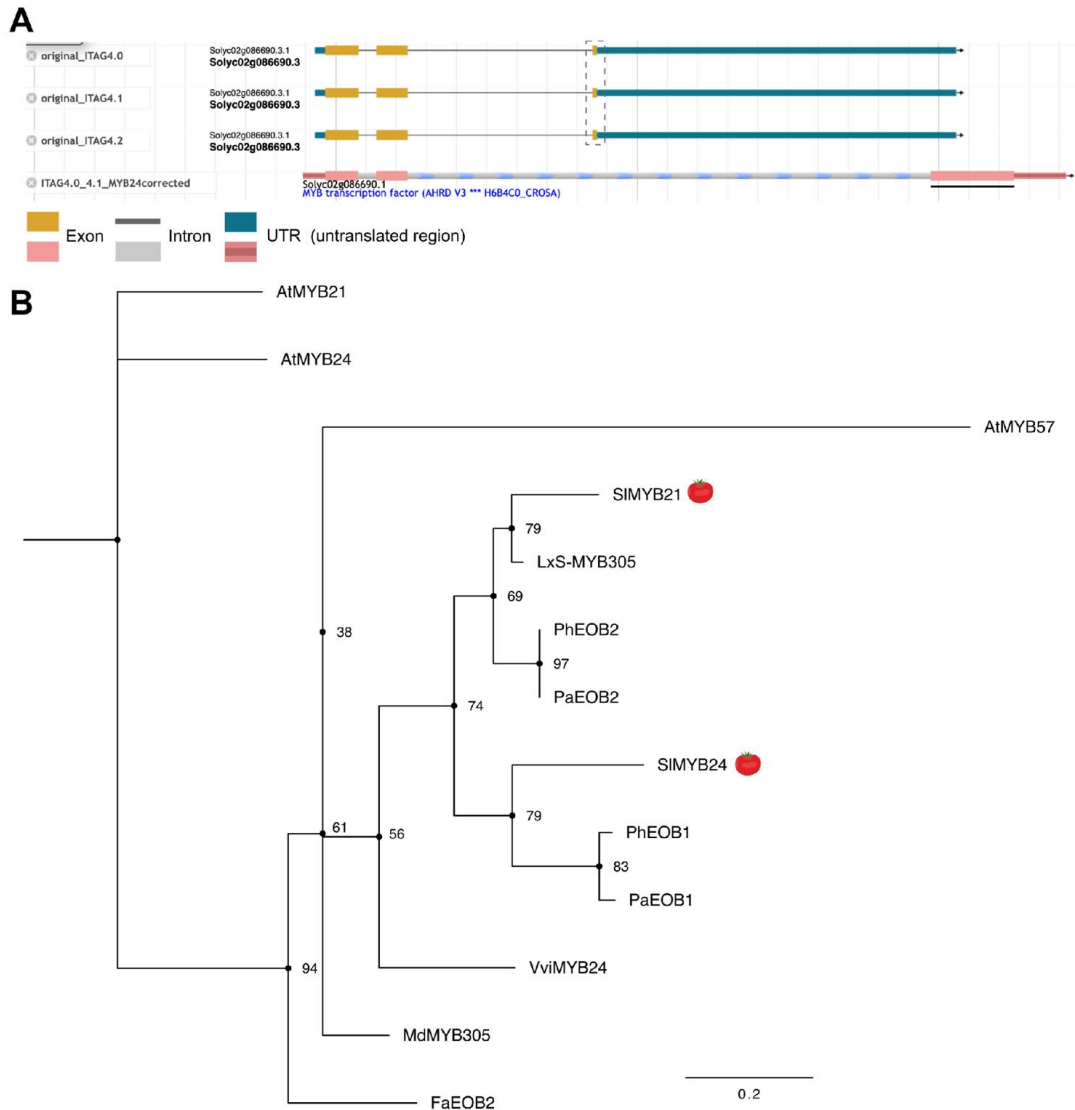

**Supplementary Figure S1: SIMYB24 is the ortholog of AtMYB24**

A) The structure of *SlMYB24* sequences in ITAG annotation version. The dashed box indicates wrong exon position and the black solid line shows the correct third exon of *SIMYB24*.

B) Phylogenetic analysis of S19-MYB proteins. The gene or protein IDs are as following: AtMYB21 (AT3G27810), AtMYB24 (AT5G40350), AtMYB57 (AT3G01530), LxS-MYB305 (EU111679), AmMYB305 (P81391), PhEOB1(AGC00814.1), PhEOB2 (EU360893), PaEOB1 (Peaxi162Scf00129g01231.1), PaEOB2 (Peaxi162Scf00080g00064.1), MdMYB305 (MDP0000344978), FaEOB2 (KM099230), and VviMYB24 (NP\_001268062). (supports Figure 1A)

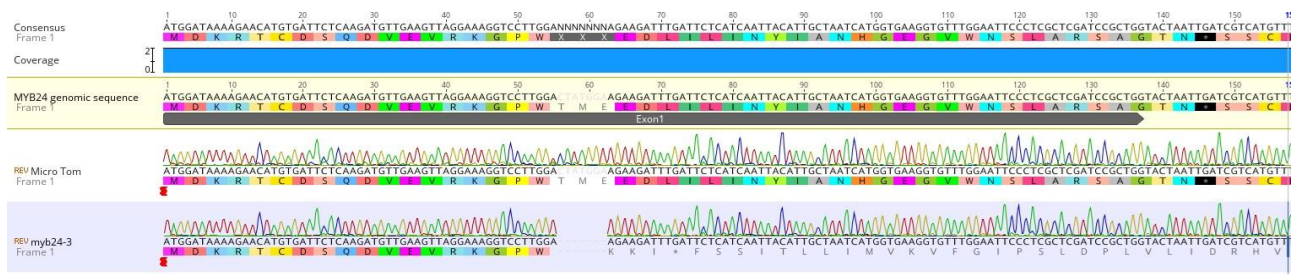

**Supplementary Figure S2:** Sequence alignment of *Slmyb24-3* to genomic *SlMYB24* sequence unveil a premature stop codon. (supports Figure 1E-G)

**A**

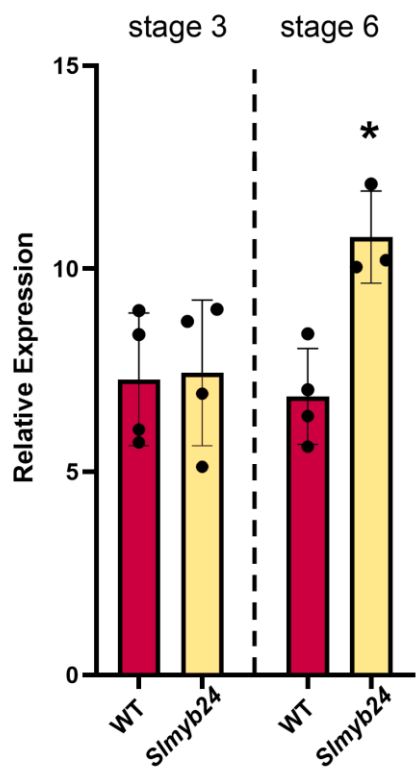

**B**

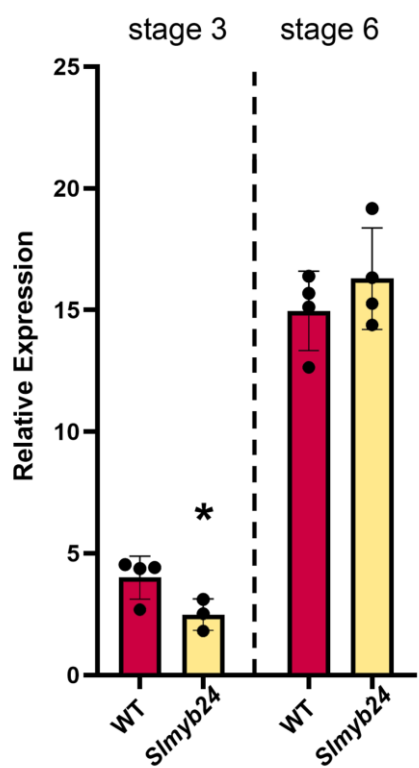

**Supplementary Figure S3:** *SlMYB21* expression shows only minor changes in carpels and petals of *Slmyb24* in comparison to wild type.

A) *SIMYB21* transcript levels in carpels.

B) *SIMYB21* transcript levels in petals.

Transcript accumulation was determined by RT-qPCR in relation to *TIP41*.

Shown are means from 4 biological replicates indicated by dots. Statistical differences between wild type and *Slmyb24* were tested using Student's t-test (\* $P \leq 0.05$).

(supports Figure 1E-G)

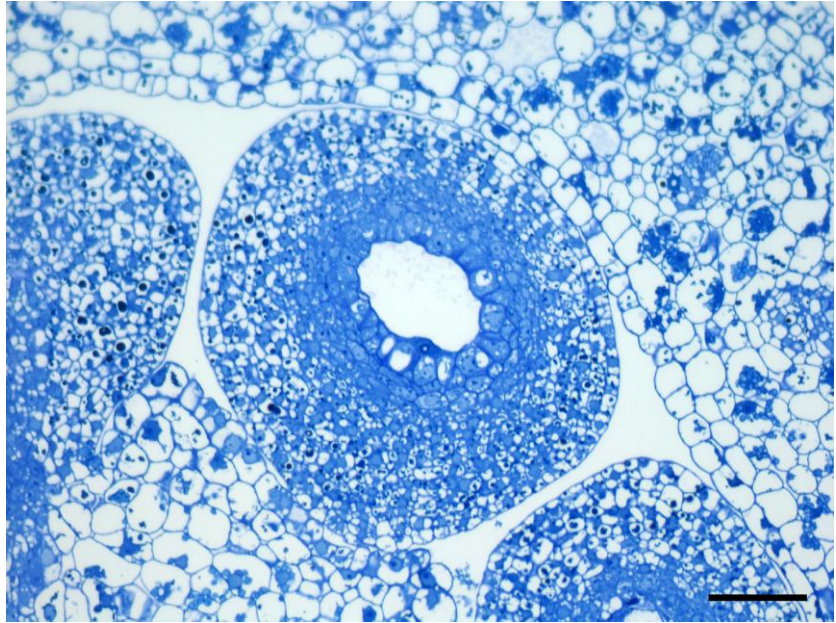

**Supplementary Figure S4:** Ovules of *Slmyb21 Slmyb24* at stage 6 show altered morphology and early PCD in the innermost cell layer of the nucellus. Compare with Figure 3 in Schubert et al. (2019). Detail of a semi-thin cross section of *Slmyb21 Slmyb24* ovary stained with toluidine blue. Scale bar = 50 $\mu\text{m}$ (supports Figure 2D)

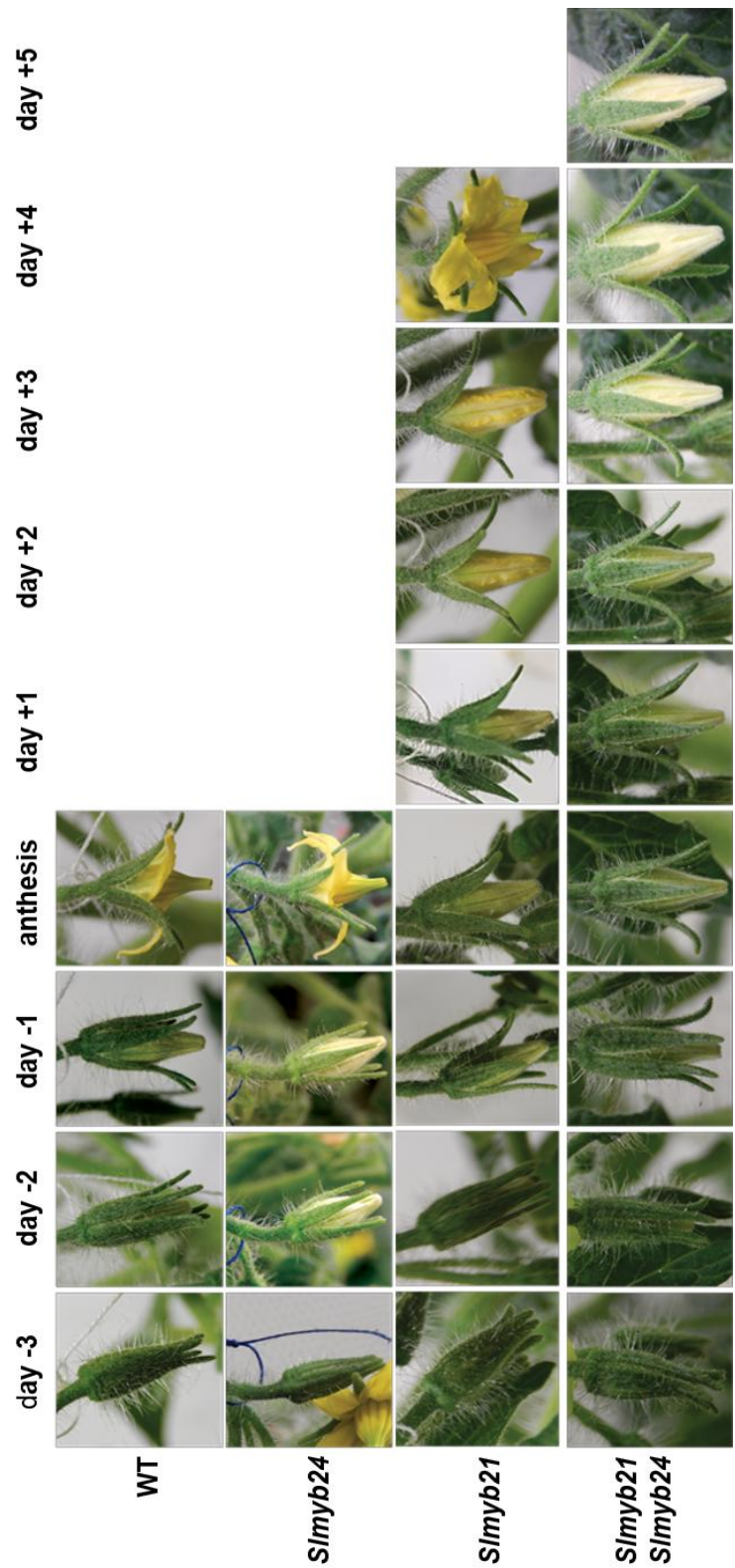

**Supplementary Figure S5:** *Slmyb21*, *Slmyb21 Slmyb24* but not *Slmyb24* exhibit delayed flower development. (supports Figure 2D)

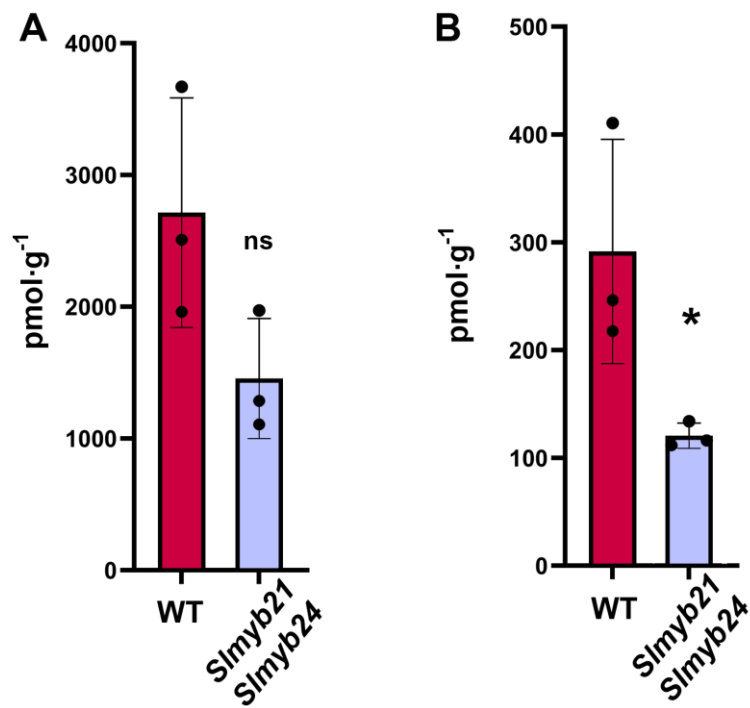

**Supplementary Figure S6:** Jasmonate levels in petals.

A) JA,

B) JA-Ile.

Petals at stage 3 were extracted, and contents of JA and JA-Ile were determined. Mean values ±SD are shown ($n \geq 3$ independent pools of petals). Data were compared between *Slmyb21 Slmyb24* and wild type by Student's *t* test (\* $P \leq 0.05$).

(supports Figure 2F)

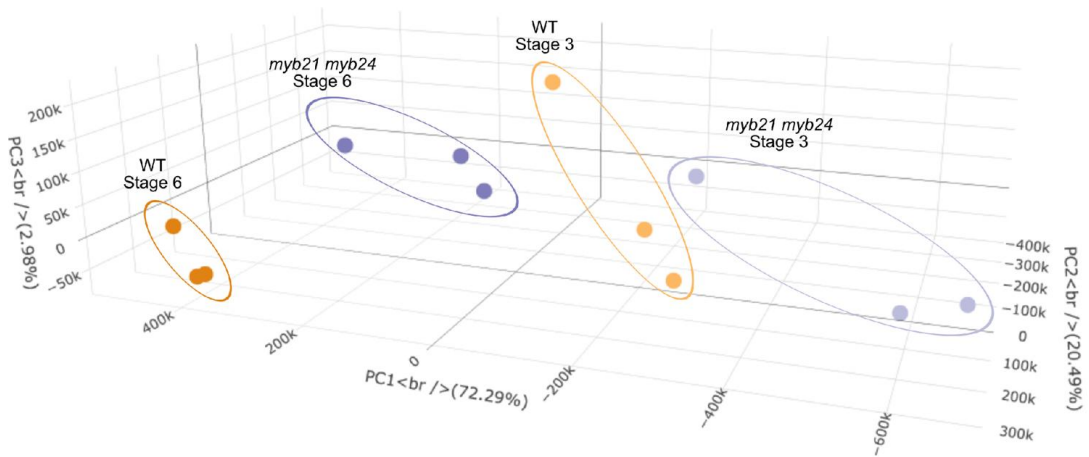

**Supplementary Figure S7:** Principal component analysis (PCA) of transcriptomic data from petals of WT and *Slmyb21 Slmyb24* double mutants at developmental stage 3 and 6. PCA shows clustering of samples by genotype and developmental stages, with PC1 accounting for 72.29% of the variance. (supports Figure 3)

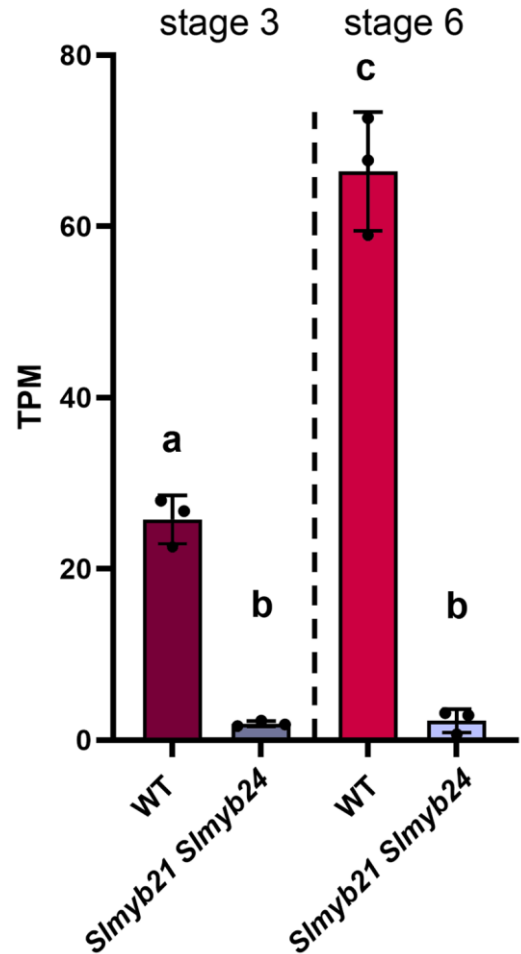

**Supplementary Figure S8:** *LIN5* transcript levels are impaired in petals from *Slmyb21 Slmyb24*.

Transcript accumulation of *LIN5* in petals was determined from RNA-seq data and is given as TPM.

Shown are means from 3 biological replicates indicated by dots. Different letters show statistically different values according to ANOVA followed by Tukey's HSD ($P \leq 0.05$).

(supports Figure 6B)

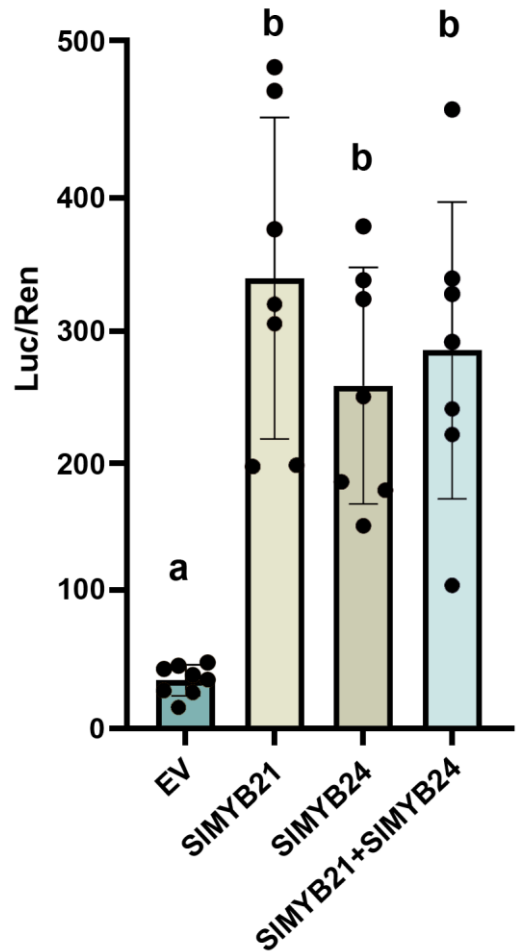

**Supplementary Figure S9:** *uprLIN5* is activated by SlMYB21 and SIMYB24.

Trans-activation assay of the upstream region of *LIN5*. The y-axis represents fold changes of LUC to Renilla (REN) luminescence due to presence of SlMYB21 and SlMYB24 either alone or in combination in comparison to the empty vector (EV) control.

Shown are means from 7 biological replicates indicated by dots. Different letters show statistically different values according to ANOVA followed by Tukeys HSD ($P \leq 0.05$).

(supports Figure 6B)

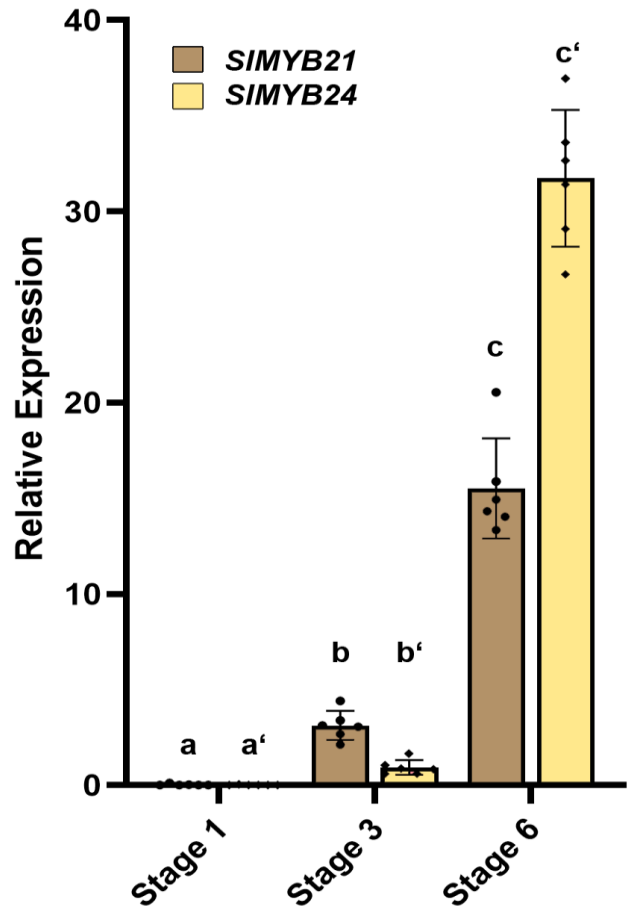

**Supplementary Figure S10:** *SlMYB24* expression follows the *SlMYB21* expression during petal development.

Transcript accumulation was determined by RT-qPCR in relation to *TIP41*. Shown are means from 5-6 biological replicates indicated by dots. Different letters show statistically different values according to ANOVA followed by Tukeys HSD ($P \leq 0.05$). (supports Figure 6C-E)

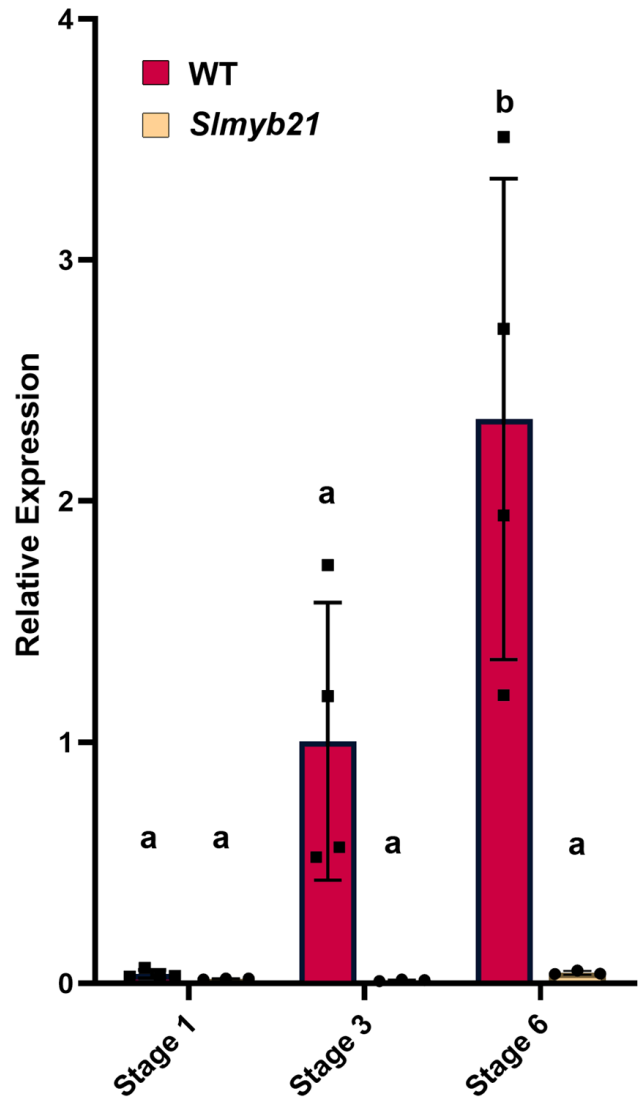

**Supplementary Figure S11:** *SlMYB24* expression is almost not detectable in the *Slmyb21* mutant.

Transcript accumulation was determined by RT-qPCR in relation to *TIP41*. Shown are means from 4 biological replicates indicated by dots. Different letters show statistically different values according to ANOVA followed by Tukeys HSD ($P \leq 0.05$). (supports Figure 6C-E)

**Supplementary data S1:** Sequences of the upr regions of selected tomato genes

***uprPSY2 (Solyc02g081330) SL4.0ch02:43319071..43322019 forward strand***

AGAATAAATATTATTTTTTTCCAAAGCAAGTACAAACAGCTACGTGTCAAATGTTGAGCCCG
CAGTTAGGCTCAACTCAACTTGTTGTGGATAGCACACGCATACGTAGGTGGGCTTTTCTGG
CCCATGCATAAATTTCCATTTTCCAGAATCAAGTTTTTACGTGGTCCGCTCTTATTTAATGTT
GACTTCATTGATTATTCTAAAAAAAATATTTCATCACCAAGCAGACCTTTCTAAAAAAAATTTA
AGATGTCTTAAAATATAACTTTGGTCACTCAATTAGTCTTAAAGTTATAAATTTGTGTCATTGA
GTTATGCCTTAGTTATAATTTGTCACTATACATAAGTATTTCGGCGTAGTTTAGTACATTACC
GAATTTAATTTCTTTTGTATTTGAACACATAGTTTATATCGTTCCACTTATGTTTGGAACATAA
TTCGTGTCATAAAATTTATGCTCAAACATATATATGTTTAAACTTATATCCTAAATTTTCTCGC
AATTTTTTTTAAAGAAATATTTCAATGAACTTCGTTTTCATTTTAAGTCAAAATAAATTAATGAA
AGAAATATTTACGCTCTCAAACATAAAATTAAAGATGTACACCCCGAAATAGAGATGTTTTGC
GCGCAAATGACAAATTTAGACAATTAGGTTCGTACGTTCTGGTGTATGACCCACATGAGTGG
GACCCACAGTCCACTTGATTAGGAGTAGGCACTTTCTAGTCCGTGGGCCCGCTGCGACTCA
TAGTTGGGTTAAAAGGTAAGAATTTTGATCCAATGAAAAATTGCCACATGTACATCGTAACC
CAGCTGCCCACTTTTATTTTGATAACGCTATCTTCCACTGATTGGGCTGTGATTTACACGTG
TCCTCTTTATACACCATCTTCTTACCCCATTACTAAAGTTGGTATTATACTTATAAATGGACT
CAATAATGCTCATATTCTTTACAATTACAGAAGAGAATAGCATTAAATTTTAGGGAACAATCC
TACCACGGGTCAAAATACAAAATAAAAATAAAATTGTTTCCCAAGTCAAAATTACTTTTTTAG
CCTACAATATTTCAAATAAATATTCTTGATTATGAAAATTGGGACTATTCATTAATGTATTTGT
AGTTTTAAATTACAATATGTGGATTTTATTGATAACCAAATTAAGAGACTTGAGCACAACATT
GCACTCTAAACTTTATCATTTTATTGATAATCAAGAAATTATTGACTATCTACCATAGGGTCC
AACCTCTATTCCGCGCCACTTATTATTTGGAAACTTTTAATGTAGCTACAACTTGTTAGACTC
ATGTGTGCTCTTTTATGTATATAATCAATTCCCTCGGAAGCTAGTTAGCTCATGTAGCACAAA
AGTGGAAATTAGGTCAATTTTCTGTATCTGAAATTAATTCATACTTTAAAAGTTAATTTAATGG
AGTAGGCTCGCGTAAGCTTTATAATAAGTCCATTGATCTCATTAATCATCAATATAGACTTAT
GTCACTTCACACGATTATTTTGATTTAAGAAACCTCAACGTTCAGTGAGATGAAATTTGACG
CTGATATCATATGAAATTAGGTCTTGAACCTAACTTACATCCTAAAAGATAACTCTAATTAGG
AGGATTATCAAGTCATATAAGAACTACCAATGCAGACTTTTGTCATTCTCTAACACTATACTT
AATTAACCAAATTTTTAATACTTTCACTTTAATATAATGTTACATTTTGTCAAGTATATGTAATG
CTAGACAAAATGTAAAATAAAATAATGTAATGCATACTCTTACAAAAAAAAAAAAAAAACTCA
AAAATCCTACTTGAATTATTTCAGATTCGCATAACATAAGACCCATTCAAGAAAATCTCTCCC
TGTCAAGAGTTATTCCATATCATGTCAATAAGAAAAACTTATTGACTCATGAAAAAATTCGCT
GCACCATATAATTATGAAGAAATAATTTCATCCACTGCACCATATCTTTCTTAATAAAACATG
GTACGTTAGTGGGACCAAACTGTAAAATTTGAGTGACAAAAGGTTAAAAGGTGGAAACTAG
GACGAGGTGGAGGAGATAGAATCAGAATCACTCAATTCTAATCAGCACGAGTAAACAAAAA
ACAAAAAAAGATTGAATGGTCTAAAGTATTCAAAAATATATAGGAGTAATTTGAACAAGAAAA
TGCTGCTAAAATTCTTGTGATCAGCAACAAGCAACAAGGAGGCCATTTTTTTTTGTTTTAACT
TTCTTTCAACTGATGTTGATGTGTTAAAAAAAAAACCTTGTTCAACTAATAGGAGCTTTCTTT
CTATTGCTATTCGCAGGTTATTTCTTTAAATTCCCCCGTACCTTTCATTATTAATCTGTTTTTT
TTTTAACTTAATATAATATTAAAGATTTGAACACCCTTTTCCTTTTCTAGATAGTCTCTACGTG
TATCAAAGGTAGTAAGGGAACTTATTATAATCTAATTGACTATATATAGTAAGCTCCAATCCA
ATTACTTGAAGAAAGTGTGAAAAAGCATATTCATTTGGGGTGAGTTGAGTAATTGGCTTTGT
GCTGTTGCTGATTTGAATTCTTGGTATTGGGATACTTTTTTTAATTTGTTGATAAAGTTTCAAT
CTTTATGATTTTTGTGTGGTCATAACTGTATCTCATTGTGTTTATTTGGTTTATCTCATTGTGC
AGAAACTCAGAAATACAAGGTTTTTGGTTGTTGTTGATGAGTGAGTGCATTATACATCTGTTT
GTGTAAGCCAAAGTCGGTTAACTTTCTTATTATGCGATTTTTATAATCGTTGGTATTGTTTGA
AATTAGTGGATAGACTCTAGTGGGTATCTACTAGGCTGAGGTGACAAGGTAACATAAAGGA
AAGACAAAAACTTGG

***uprLYC-B (Solyc06g074240) SL4.0ch06:43564023..43565230 reverse strand***

TCCATTCTCTGAAAAGGAGCACCATATTTGCCGCACTGTGGTTCATATTTCCAAGTACATTT
AGATGAACTATATCATCAGATTGAAAGGTTATTGTATAATCAATCCAGTGGATTCTCGTTCTG
GCACCTTTAGAAGTACATGTGCGGAAAAGAATGATAAGGTTTGTATTGTTGTTGACAAAGCC
TGTTGCCTTTCTCATTTGTAAATGTTCTGAACGACTCCTAAATTACTCTTAAGGTGTAAGGTC
TTCCGTGCCTGTTTGTAAATATAATGCTGTGCCGTGACTTACCTTTTGTACCATTTGTTCAAA
TGTATGGCCTGGACACCAGGGTTGTCAAAAATGTCTCATGCCCGTTTTATTGGTCTGAGAAT
GGCGTGATGCCAAATTCTGCCGCTCCACAGTGAGCATTTCGATCTACTGGAAATTGACCAA
CTTATTTTATCACTTGATAACTAAACAAAATCCTATTAACTTTAATCATACATTGTATTTATAC
CGAAAAAGTTATGCATAACTCAGTAAATTACCTTTTTTAGCAGTCAAATTCTAGATCAGTTTC
TAATTTATCAAAATGGCTTTTATAGGGTCCCAGTTCCACTAATATACCTGCCGTCCATGCAC
TGACTACAAGACAAATACCTCACTATGTTTGTTAGTGCTTGGTAATATAAAACCTTTTCTTTT
ATGAGAAAGTTCACCGAGAATAATTTTCTATTTGTGGCATAACTAGTATATAGTGCAGATTGA
CAAGAATTTAATTTTGCAGTTGGGCACATGAACAATTTTCCTCAAAGTTGTAGAAAGTACTTT
TCATTTTCTTGTCACCGAAAATTATTTATAATTGAAATTAAAACCGAATGAGCTGCAAGATTC
AAGTCGAATTTCAAAAGAATTGACCAAGAAAAAATTCAAAAATATCCCCCACCCCCTACCAA
ACACATCCTAAAGTGAGGTATAGACTGGGACTGGGATTGGGAAAAGGGTAAAATGCTTTCA
CTAGCTTAGCAAAGATTCCACTTTGTTAGCTATCTTTCTTTCTCATTTCCTTTTTTCTTTTTCT
TTTTTTTGTTATATAAGCCAAAGTAGGTACCCAAAAGCATCAATATTTTGTATTGCTTGGTGA
TTCCTCTGTAGTCCAGTATTTCATTTTCTACAAGTTCCACCTCCCTCCATAATTAACCATTAT
CAATCTTATACATTCTCTATAG

***uprNCED1 (Solyc07g056570) SL4.0ch07:64280348..64283602 reverse strand***

TGACTTTCATGTTTATGTCATATTTATTATGTGCTTCTATAATATCATTTTGCTAATATAAACAA
AACGGATATTAAGATAAATGAGATTGCTAAGGAGAAGTACAATTGTTATTTATTTTCACGCTAA
ACTCATCATTTATTTATAGATAATAAAGATAAGAATACAAGTGGTATATATTTTTATTTTATTTG
TGAGTTTTGTTATTAAACGTAAATATTTCTATTTACTATATTATTTTTATAGTTTGATATAAATAT
TCAGTATAAATAATACTTCGTTAACCAGAAAAAATAGGATATTTCGCTTTATTTGTGCTTTTTAT
AACTAAGTTATCCTGTTTAATGTAAACATTGAATACAAGTGATATTTCGTCATTAAGTTCTAAA
AGGAAAAAGAATAAACACGACGTAATTTTTGTATAATTTATATCCAAATCAAACTTAACTACCT
AAGTAGTGCTTTTTCCATTTTCTAAAACTCTTTTTTTTTTCATTCAATATCTGATAATTTTATTGG
AATCTGATATTTCAAAGTTCATTCTGCAAGATATCGTTCTCAATAATGCTTTTTCTATTTTCAAA
TACTTAAATTAGAAATGTTTTATTAATAAGTAAACATTTCACGTAATAACTGATATTGGTCAACT
TTTCTTTTCACATCAAATGTTTATCTTTGACCAATAAGTTTGTTAAAGAAAATAATAATGCTAAT
AACTTTATCTTTTTTTTTTCCTTCTATTTGTTAATCTTTCGGTGTTAAATTTCTTTGATACTCCCT
ATTTTAGCATAAACTTTTGACTTATTTGAAAAGACTTGTATCTTTTTGTAGCTCTCTAGCATATA
ATTATGGATTAGATATACTTACATTAAATTAAATATTTAAACTATAAACTCTTGTCATATATATT
CAATATAATTTATTTTAATTTCCTCCTTTTTTAAGTGTTGAGAATTGAGAGTTGGATAACAAGAA
TATTCCACTAGTCAACTTTTGATTTTGATGAAAATTATTATTATTGTTACATTATATGTACAAAG
TAGATCTAAATAAAAACTAGAAAATTCACAATAAAGGATGAAATATTTAAAAAAATTCAAACAA
CAAATATGATGTAAGAGAAACATATACATGTGTCAAAAAAAATTCAGAAATCAATAAATTATAT
ACTTCACACATCTACTATAGTATCGTAATTAATAAGTTATATACTTCACACATCTACTATAGTAT
CATTAATACAAATGAGCTTGACAAGTCCTTTCTAAGTATCAATGGTGGAGATATTCTTTTATAA
ACGAATATAATTAAACATCGTTTGACTATTCTATTGATAAAATCATATATATGAAACTTCATTTT
TTAAAGGAGAGTGAACTAAAGTTATTGCTTTAAAGTTTAAAATAATACAATGTAGACATTAATT
TAGAATAAAAGTTTTAAATTTTGCAACATTAAACGCTAAACTCAAAAGTATATACAAGTTGAAT
TGTAATAATAAGCCAATTTCACTCGAAAAATATTGTATACAAAATTAAATATATTTTTCAAATTT
TTTACTATTAATTTGTAGGAAAGAAGAAAATAAAAACACACGTTTACAACATAAGATGATAATC
CCTCAATCCCTCTAGTTCACATTGAGTAGATTATTTTTCAGAGTTCAAAATAAATCAATTTCTT
TTATATGTCTACGTAATTATTTAAAAAGAATTTATCATCCTTCTTTATGGAGTTTTTAACTATTTT
AAATTTACTACTAGAATAATTTAAAAATAATAAGAAGGATCATAATGAAAAATAACCTTCCTGA
AATACTTTTTTTTTTCAATGAATGTACCCCCAAATCCCAACAATCCACTTAACATGAATTAAAT
GGGGTGCTAATTAGGATGGAGCCTCTCTCACCCCATATATTTTTCTATTTTTTCGATATGAATT
GTGATGAAATGATTAAAGACTCTTTCATTCTTAATTATCTTTCAATTTAAGCTTTACATATGAAA
AAAAACAAATATTCAAAATGTCATTTTCAAAAATAAATTTTACTATATGCAAATTCAAATTAATC
GTAATAGACGCTCAGTGATCGATCATAAACACAGAGAGTTAGGTAGTATTGTCAAATTTATTT
TCATAACAAAAAATGTAGGGACACCCACCCCTTCTCCCCACACCCACATGGGTCATTTGTTG
TATTAGTAGTAATATATAGTAGTAGTAAAGAGTTTAATCCTCCAAATATGCATTTTTCACATATC
CAACACGTGGTCTTCTCTTACATGTTAGAAGTGGACCATAATTCCACACTCTCCCCAACTCTA
TATATACAAATCCTCCCCTTCTCTCTTCTATAAACTCAACTCAAAAGGAAACTAAACATAAACT
CTCCTCATTTCCCACCTCTTTTCTCTCTACAAAAAAAAAAAAAAAATCCCCTTTCCACAACACC
AAATTGTACTATTAACATCCAAAAAAAAGAAAAAGAAAATAGGTAGCT

***uprTIP1.1* (Solyc06g074820) SL4.0ch06:44020005..44022623 forward strand**

TAATAATTAAATTAATAGGGTGGTTATGTATTATGAAGGAATCCATGACTTAAAGGGAGATT
AATTAATTAGTACCTACTACCAAAGTTTACAATCAACAATATTTGTGGATTCCTTTGGTTATT
TATGTAATTTTATTTCAATTTGGATTATCTATCAGAAAATTAATTAATTTCGATTATTAAAAAT
GTACTTTTGTCTACTTAATAATAAAATATATAAAATTTGTTTTGCTCCATCAACTAAGAAGGA
GATGGACGTAACAACTATGTGGACCACTTGAATTACATCTTTTAATTTTTTATTCTTTCATAT
GTTGTTATATAATAAAAGTTCATAAAAGAGGACCAAATTTATAACTATACGTATTTATCGGC
ATCTTTCGGATCGAAACAAATTTAATGAGCCTGCACGTATGTCAATTTTTTTTGAAATTATTT
GCTTTTCGAAGAGTCTTCTAACATGTTATTATTGTTATTGTTAATATCACTTTAATCGGGATA
GTTTATTTTAACATGCTATTGTTAACAATAACAACTATAAAGGATTCCTATCCTCTTATTATT
CTTATTTTATAAAGATAAATATTTTCTTTAAAAAATTAACAACTCAAATAAAATATTGAAATGT
ATATAATATAAATCATTATATTTTAAAATTCAAATTTCAAAATTCACTTTTACAATCAAATCGT
GACAAAAGGTCAAAATCAACTACAGTACATATACTTATTTTAAAAATAAAAATAATAATTACA
GCTTAGTTGTCAAAGCTATGATGTATATAGAGTATTATCAATTAAAAGAACAGTATAATGAA
TGTACATCTTTCTGGAGGGAAATATAATATGCACGTTTTTCAGATCAATAATTAAATTATTG
GCCAAATAGAAAAGGCCGGCAAAACCTTATCAGAAATTCCTCTTCAAACTGATATTTAAATT
ATTAAACCCCACTCTAGAGTGATTATTCCTTTTTCTTTTTATTTGATATTTATATTAAAATTTT
AATAAATATAAAAATTTATTTGGAATAAAATTTATTGATCTTTTATTAATTCTCTTATAATTTTT
TTTATATCTTAAAGTTTGAAGTCGAAATTACTAATAAAAAAATTAACATGATTAGAGTCATTA
TCTTCATTATTATACCAAAAGATATCATATCTCTAGACCCTATACCTTTTTTGTCATTTCTTT
CCATCGACCTAATCGATTCCCAATTTTTTAAAATTACAACTTTTTGCATTTAAGGAAAAATAT
GAAATGTAGCCTTTATTGCAATCATTTACAGTTGTTCACGTACACCACAAAAACTTCAAAAA
CAATTAGAAGTTCGAAAATCCCGGTATAAGTTGTTACTTTTATTATCTTAATTTATGTGATTT
TTTTTGTCAATCTCAAAAATGATAAGCTTTTATATTAAATAACAATTTAATTTTAAAATATCTA
TATTATTCATAATGAAATATCCATCTAAATTTCTATCATTCACTTGAGACCACAAATTTTAAA
GGTTTTTCTCGATAGATATTTTCTTTAAAAAAATTATAAATATTAGATTCCAGTCTCAATCAA
GACAAAAATATTAAGGGTTGCTTTTTATTTAGTTAGGCCTTGATAAATATAGTTATGATAGA
ATTGATAATTATACTTCGTAAAATGAATTAAAATATATATAAACCAACTCGTACAGGAGATTT
ATACATATGAAAGTTGTGATTTGGCCGGAAAAATAAATGATATTTAAGAGAAAAATCTAGAA
TGGAAGATAAAAGAAAAGATTGGGAACCCTAGCTAGTTCTAGATCACTCCATGGTGCATTA
TTGCTATTTACTAACTATTTCTTTTACCACTTTCTTCCTTATATAAATTTTACGCTCTTTTATT
GCAGTCATTTAAAATTATACACGTAACTTCAAATCTAATATGATATAATGAATAAGACTGCT
TCTCGTTTAATCAAAGATGTTCAAAATTTTAATTGGAAATAGTGATCTACCCAAATGGGATC
TATCCTACTGATTAATAAGACTCTAATACATATAATAATAATATAAAAAAAAAGTACATTTAT
CTCCGTTTTTTTTTAAAAAAAAAATAGTTTCATTACAAAAGTTAGTGATTAACCTAATCAATA
ATTGCCATTTAAAAAACAGATTAAAAATAATTAAGAGCCCTAGTTCCAGATCATTCCATGGT
GCATGATTGCTCCGACCGAAAGCAATAACAAGGGGTGGGTGGGGTGGGGAAGAAGGGG
GGTATGGGGTATACAGTACTCCCACTATTTCATGTCTTCATCAAACAAAACCTAAATGACG
TGGCTCATTGTCGTTATTTCCACGTGTCATCACCACATGAAATTGGAGGTGAATTCTTAGC
TGTCAATGCAGGCCTTTCGACCTCTACGGGACCGCTCGCCCACAAATTTTGGTTTGAAAT
CTTATTACATTATGGTCCAAATATTAACCGGTTACCTCCTATACCCGTTACCTAACCCCTCA
TGCAATTTACAAATTTCACAATATAAATCCATGCAAAACTTTCCCTAATTTCATCACTCCCC
AACTTGTGCCCCCCTTAATAGTGAAAATTTTCAATTGTTAATTTTCTCGTTAAAA

***uprMYB24* (Solyc02g086690) SL4.0ch02:47353665..47356323 forward strand**

GCATTATGCGTGTCTGTACATTGAACTAGGAGGAGTATGATAGTTCAAAC TAATTAAAGGGAATCA
CATGTAATTAACATGCTCCTTTTGCCTAACTTCTCTATTTAACTTTAATTTTGCACGATTCTCTTTAA
TTTAATTAATTTCTTAACGAGGAAGAGGATTCTTACTTGGATACGAATTAGTAACGGCAAAATTAAC
TCATATTTTGGTAACCCGCCCAATTCACTCATTTATTAATAGCTTGGGTGATTGATTTTTTTGGATG
GGTAAAATATGAATACATACCCGTATTTAATCCATAAAAATATGAATATTCATGGGTTAGCCTAATG
GATTTAACTCTCAACTTTTATTCACCCAAAATTTATAATATCTGTAAAAATGTTGTAAAATAATGTCT
CTCTCTCTCATTTTTTTTTTTAATTTCAATGTTCTTATTATTCTTCTCTATAACTGAAGAGGTAAAAGT
CCAAAAAAGTATATACTATATTACATGTAAAGTAAGTGAATCATGTATTAGTATTGATATTGTGTAAA
GTTCATCAAGCACTCTTTTACGTCTAAAATGGAAATGTTCATTCGCTTTAAATTAAAAATATTTTTCC
TTGTTATTACATAAATTTATTTTATATCTAAAAGTTATATGATTTTTTTGTTGACTTTGATTATACAATA
AAACTAAATAAAGTATTTTTAACTTTATTCATCTAAGAAAAAAAAACATTTTATCCATGTTTAATTCTA
AAATAAAGTAATTTACTCATGAAAATATGAATAGACTTGTATTTTACCCAATCTATTTGTTATTCAAC
TCATTTTCACCCATATAAAAATAGGCGGGTTGAATTATTATCCATTTTATATTGATTCATTTTCAATC
TGTCCAATTAATCCAACCCATTTATTTACCACCATTAATACGAGCCAACAATAATAACATATTCAATA
TAATTTCATCACTGAATTTGAAGAGAATTGTGTTTAGGCAGACTTTTATTTTTATTTTGCAAAGGTAG
AGACAAACCCATAAACCAATTTTTTCTCTTTGGCTATATATATTATTATTATTACTACTACTACTACT
ACTATTATTATTATTATTATTATTATTATTATTATTATTATTATTATTATTATTATATGGAAAAAGCTAT
TATATATGTGAATACGTCCGGTAGATGTGCAATTTGCGTGCGGCATGTACGTCGTGCAATAGTTTC
ATTAATTCAGATATTAGTATATAAATCTATTGAATTTAATTAACCTTTATATGGACTAATTTAATCACA
AGTTAATAAACCGTAGCAAATATTAATTAGTAAATATAACATAGTGTTAAAGTTCTATGCATAGATAT
AGAGATCACCATCGATATATAAAAGGTCAACTTTCACATATTGCAAATAAAAAATTCATATTTGTAT
GCTATAGCAAAGTTTGCATAATTGCGCTCCATAGCAAACACAAAACTTTATAATTCGCTATATATAT
TCAATTGTATAATTCGCTGGCCTAAATTATATAATTCGCTGGCCTATTTTCGTTGCAATTGTATAATT
CGCTATCCTATTTCACTACAATTGTATAATGCACAATTGTATAATTCGCTGCCTATTTCGCTGCAAT
ATTTTTATAAAATTTACTTTGCATACAATTGAATCGAATTAAAATATATGTATATTGCATAATTATAAG
TGTATAGCAAGAAAATATATGTTTTTCTCTCGCTTTATACAAAAACAGAAACACAATATATACACTTC
TGTTGTATAAAGCTAGAGAAAATTGTATTTCACTGCAATTGTATAATTCGTTGACCTTTTTCTCTGCA
ATATTTGTAAAATTTGCATTTGTCTACAATTGAATTGAAGTATATGTTTTTGCATGTGTATATACAAT
TTTCTCTCGCTTTATACAAAACAGAAACATAAATTATACACTTCTGTGTATAAAGCGAGAGAGGCGA
GCAGAGGGAGAGTGACGAGCGAGAATGGGAGAGTGACGAGTGAGATTTTTGGGAGAGAGGCGC
CTGGCAAATTTTGGCTAACGTTTGCTATGGAGCACAATTAAATCAAACTCTAGCTACTCCATTTATT
TTAGATTATTAGTTTGGTATTATATACAATTTTCCCTATATAAAAATATTGTGCGATTAAATAACTAAA
AGGTACATATAAAGTTTTATACATTGAAAATACGTAAAATATATACAAAATAATTCATTGATAATGTA
AAAAAAAGTTACACTTCAATTTAAATCTAATGTGAAATTATTGGGGTTTCTATGCATAAATAGCTAGT
TTCCTTGTTTACTTATTAAAAGCAACACTAATATTATACACTTTAAATAAATTTAACTGTCAGCGTAT
ATAATTGATCAAATCGAACTAATTAATTAGTAAGCACTAGCTCGTTGTGACCATGCACGCACACAT
CGTTGGCTGTTATTTATGCTCGTCAATTAACGATAACGACGATTAAGTATAAGTAAGAACTTTATAG
TATATAAATTATTTAGCGCAAGTTAGGTTAAAAGTTAAAAGACTTAATTACTGTTTTAAAATCTCATT
TTAGTTGTCCCCTATAAATATCAGCCCCAACTAACGTTAGTTCCTCGTCAAAATCTCCAACTACAAA
AAAAAAAAAAACTTTTTTCTTCACATCCCATTCCTCTTCTCTTCTTTATTATATAAAAAAA

***uprLIN5* (Solyc09g010080) SL4.0ch09 SL4.0ch09:3505656..3508224 forward strand**

TGAGTAAGATTCAAGGAGAAAGATAGGTGGATGGATGAAAAACAAATTCATATAATTTTGAATTGA
AAGGAAAAAGAGTCAAATTCAAAATTACTATCGAAATACTTCGATTTTGTGTCTCGTGCTAAGGGA
AATGGTGAACGAATGTCAGAAGAGGGATGGGGTGGGATGTTCTGGGAAGAAGAAGAAGAAAAG
GGGAACGAAATATTTTTTTTTAATTATTTTTTTTAGTGAAAACATAATTTTTGTTTATTTTATTTTTTT
AATTTATGTTTTAAGTGTGTTTATGGGCCCAATGCCACATGTCTCTTTTTTATTAGCTATTATTTCA
CGTTAGCAGCGAGTGTAGGACATTCTCTCTTTACTTTTAATTAATTGTCAAAATATATATATATCTC
TTTAAAATACACTATTAAATAATACATCGATAAAATATCTTTTCCAAAATTTGATTATTATTACAATA
ATTTCGATCAAAATTCATGAGGTGTTTTTTCTTTTTTTATATACAATTCACTAATATTTGGCAATGGA
ATCCACTAAGAAAATCAAAGGTGGAAAAACTATTAACTGCAAATGTTGCAAGACATATTTATGCAA
ATGAAAAGTAGAAAGCTAATACTTTTTGCCAAATTCATCAAAAAAATGTAATGTGTAAAATGCACTT
TATTGTTGTTATGCATATATGTTCATCACCTATTCATCTATCCATTATATTTCATTTTCAATAATTAC
TGTGGTCCTATTCATTTTATTTTGGAACTCAAATATCATGATTACATATTATTTAAAATTCGCGTTA
CGATGAAATATAGGCGAAAAAAACTATTAATCTTTCTCATGAGGTGAGGTGAGGCGAGAAAAACA
AAACGTTATATATTATTTGATGTCTTGTGTGAGGTGTCACTATTCTAGTTGTGTTTTATCTCCGTCT
ATACTGAAGTTTAAAAGTACATATATAGCTAATCGAAAACATATTTTTATCATTGAAAATTACAAAT
TTTTTATGTACGTCTTTGTTGAAACGTCGATAAAAATATAAATATTAATCGGAAGTCGGTCAAAAAT
ATTTCTGACAAAACAATTCTATTCCATCGATAATTTTTACTCTTTTTTTTTTTTAAGTATTGATCAAT
TTATTCATCTAGAAGGATAAAGTCTACATTTTTATTTGTTTTAAACATATATTTACACAAATGGTCTT
TTCAACTATATGTTATAGAATTTGTGAATTAAATCAAATATTATTTTTGTTATGATTTTAGGTTCGAG
TCGCTTATAAAAAGATTGTAGTTCATAGCCAAAATTCTTCTCAAAGGCGGAATAATAATTTTAAATT
TATGAGCGTTTTGAATTCTAAAACAGATAATTTATTTAATAAATTATATTATACATATGAGAGATAAT
TTATTCAATAAATTATTTATTTATATTATATAAGTGGAACTTTAATATAAATATACGAAACGCTTATG
CTAAAATTGATAACTTGTATTTATTTTAATTTTAATGTTTTTGGAGGGAGAAGGTAGGTAAAATTTG
AACATATATTGGTACGTGACAATTCTAAAATGTCTTGAAAGGTCGTGTCGGTGCTACGATTTTTAA
CGAGAGAGTTTAAAATTTAAAGAAGTAAAATAGATAATTTAATTATGTATAAATAATAATATTTTCTA
TCAGAGTGGATCCGAACGAATATTATTTTGAATATTATTTTGAATTAAGGCAAAACAAACACTGTA
TATATGATTAATTTTCGTCAACCAATAGTCCCTTCTCTAGTTTTCTATACGTCAAATTAGATTGTTT
AAACACAAATACTTATATGATAATCTTTTACGTGTAATCGATACATAGAAAATAACGTTATTATTTTT
TACCAAGAGTCTTATATATACTTTATTTGTAAATTATTACGTCAGGCTTTATACGTTAAATTATTTTG
TCATGTAAATTAACTTTACTTTAGTAATATGAAAAGTCTATACAAGCATCGAGTGTATAATTTGACT
TTTATTCGATTAAAAAAATACACCTTATATATACAAATTAAGATTATATTTTATATTTGTATAGTAAA
TATTGACCTATTTATTGACGATATAAAATCACCCCCTGATCGTCATGCTTGAATGAAGACACCTGA
TTGATTTGATTTTTTCACATGACCCCTCCTCGCTCTTCTCTTTTACGAGTTTGAATTTTCATATAGT
TTTAATTATCATGATAAAAATGAGTGAATATATTTTTTAAAATTATATATTGATACATATATCGAGAT
AAAAATTAAAACTAAATCTCATAAAAATGAAAATTTTGATTTCGTCACTAAACGAAAGAAAGAGAA
GTCACGCGTAAGAAAAATACGGCTAATATAATTGAAGTTTATCTTTTAAGAAACATTATTTACATCC
ATTTTATTTTAATTAAAGAAAAAGAAAATATAAAAATCAAACCGTTTTTTTTATCCACTCATCATAGT
TGGTTTTTGTCTATAAATAATCCAATATTCTTTCTTTTCTTCTTCATTATTCTTCAATATCTTTCTCC
AAAATCACAAAAA

**Supplementary Table S1:** List of primers used

| GuideRNA for GG-assembly |  |  |
| --- | --- | --- |
| MYB24<br>guideRNA1 |  | ttggtctcaattgtaggaaaggtccttggactaGTTTTAGAGCTAGAAATAGCAAG |
| MYB24<br>guideRNA2 |  | ttggtctcaattgttcccctccgaacatcaggtGTTTTAGAGCTAGAAATAGCAAG |
| Gateway cloning |  |  |
| attB1 MYB21 |  | GGGGACAAGTTTGTACAAAAAAGCAGGCTTCATGGATAAAATATGCAACTCTCAA |
| attB2 MYB21 |  | GGGGACCACTTTGTACAAGAAAGCTGGGTTTTAATCTCCATTAAGCAATTGCAT |
| attB3 MYB21 |  | GGGGCAACTTTGTATAATAAAGTTGTGATGGATAAAATATGCAACTCTCAA |
| attB4 MYB21 |  | GGGGCAACTTTGTATAGAAAAGTTGGATTAATCTCCATTAAGCAATTGCAT |
| Genotyping of mutant alleles |  |  |
| SIMYB24 | for | TTGTGACCATGCACGCAC |
|  | rev | ACACGTTGTAAATTCCCCATC |
| SIMYB21 | for | CTCTGAGCTCACTAAGTGAAC |
|  | rev | GGGACTATATGTTATGGAGTGG |
| RT-qPCR |  |  |
| NCED1<br>Solyc07g056570 | for | CATAATCGAAAACCCGGATG |
|  | rev | AACTTTTGGCCATGGTTCAG |
| PSY2<br>Solyc02g081330 | for | AGTTCTGCTAGTAGATGGCC |
|  | rev | GGGCACTAGAGATCTTGCAT |
| LCY-B<br>Solyc06g074240 | for | TGGGATGGAGACATTGTTGA |
|  | rev | AGCCATGTCCGAAAAGACAC |
| BCH2<br>Solyc03g007960 | for | ACATGTTCGTTCACGATGGA |
|  | rev | TTGTCCGAGTGATGAAGCTG |
| TIP1.1<br>Solyc06g074820 | for | TCATCACTCCCCAACTTGTGCC |
|  | rev | AAAGCCATACCAGAACCCTGACCT |
| PIP2.8<br>Solyc01g111660 | for | ATTCCCATATCCCTGTGTTGGCTCC |
|  | rev | AGCTGCAGCTCTCAAAATGTATTGG |
| MYB24<br>Solyc02g086690 | for | CTCGATCCGCTGGTTTGAAG |
|  | rev | TTTCCCCTCCGAACATCAGG |
| MYB21<br>Solyc02g067760 | for | TGCTGGTCTCAAACGTA CTG |
|  | rev | TAGCCATCGGAGTCTACAAC |
| TIP41<br>Solyc10g049850 | for | TCAGTGGGAGGATTGTAAGG |
|  | rev | GGTTCTTTAGACGCCAATGC |
| Trans-activation cloning |  |  |
| uprTIP1.1 | for | GTCGACTCTAGAGGATCTTTTAACGAGAAAATTAACAATTGA |
|  | rev | ACCATGATTACGAATTCGTAATAATTAAATTAATAGGGTGGT |
| uprLCY-B | for | GAATTCGAGCTCGGTACCCGGGGATCCATTCTCTGAAAAGGAGCACCATA |
|  | rev | GTCGACCTGCAGATGGAAGACGCCTATAGAGAATGTATAAGATTGATAATG<br>GTT |
| uprPSY2 | for | ATGATTACGAATTCGTTTTTTAAAGAAATATTTCAATG |
|  | rev | CGACTCTAGAGGATCGCTGAAACAACAGACACC |
| uprNCED1 | for | ATGATTACGAATTCGTGACTTTCATGTTTATGTCATATTT |
|  | rev | CGACTCTAGAGGATCAGCTACCTATTTTCTTTTTCTTTTT |
| *uprLIN5*<br>Soly09g010080 | for | ATGATTACGAATTCGTGAGTAAGATTCAAGGAGAAAGA |
|  | rev | CGACTCTAGAGGATCTTTTTTTTTTTGTGATTTTGGAGA |
| *uprSIMYB24* | for | GAATTCGAGCTCGGTACCCGGGGATCCGTTTGATACGTGTACTTGG |
|  | rev | GTCGACCTGCAGATGGAAGACGCCTTTTTTTATATAATAAAGAAGAGAAGA |
| *uprSIMYB24*<br>500bp | for | ATGATTACGAATTCGAACCTTTATTCATCTAAGAAAAAAAAA |
|  | rev | CGACTCTAGAGGATCTTTTTTTATATAATAAAGAAGAGAA |
| *SIMYB21* | for | GTTTTTATGAATTTTGCAGGTCGACATGGATAAAATATGCAACTCTCA |
|  | rev | CAACCAGACAACCACAACAAGCTTAATCTCCATTAAGCAATTGCA |
| *SIMYB24* | for | GTTTTTATGAATTTTGCAGGTCGACATGGATAAAGAACATGTGATTCTCAAGAT |
|  | rev | CAACCAGACAACCACAACAAGCTTAGTTGTTTGCATCATTAAGCAATTGCAT |
| **EMSA probes** |  |  |
| EMSA probe<br>labelled | for | Cy5-TTATTTAGCGCAAGTTAGGTTAAAAGTTAAAAG |
|  | rev | Cy5-CTTTTAACTTTTAACCTAACTTGCGCTAAATAA |
| EMSA probe<br>cold | for | TTATTTAGCGCAAGTTAGGTTAAAAGTTAAAAG |
|  | rev | CTTTTAACTTTTAACCTAACTTGCGCTAAATAA |
| EMSA probe<br>mutated | for | TTATTTAGCGCAAGAACGGTTAAAAGTTAAAAG |
|  | rev | CTTTTAACTTTTAACCGTTCTTGCGCTAAATAA |
| **EMSA in-fusion-cloning** |  |  |
| *SIMYB21* | for | AGGTCTCGCATGGATAAAATATGCAACTCTCAAGATGT |
|  | rev | AGGTCTCGACATTTAATCTCCATTAAGCAATTGCA |
| *SIMYB24* | for | AGGTCTCGCATGATGGATAAAAGAACATGTGACTCTCA |
|  | rev | AGGTCTCGACATTTACGAATTGTTTGCATCATTAAGCA |
| **split-Luc in-fusion-cloning** |  |  |
| *SIMYB24* | for | CGGGGGACGAGCTCGATGGATAAAAGAACATGTGACTCT |
|  | rev | ACGAGATCTGGTCGACGAATTGTTTGCATCATTAAGCA |
| *SIMYB21* | for | CGGGGGACGAGCTCGATGGATAAAATATGCAACTCTCAAG |
|  | rev | ACGAGATCTGGTCGAATCTCCATTAAGCAATTGCATTGAC |
| **Yeast-two-hybrid** |  |  |
| *SIMYB24* | for | GGGGACAAGTTTGTACAAAAAAGCAGGCTTAGATAAAAGAACATGTGATTCTCAAGATGTTG |
|  | rev | GGGGACCACTTTGTACAAGAAAGCTGGGTATTAGTTGTTTGCATCATTAAAGCAATTGCATGG |
